# Lanthanide transport, storage, and beyond: gene products and processes contributing to lanthanide and methanol metabolism in *Methylorubrum extorquens* AM1

**DOI:** 10.1101/647677

**Authors:** Paula Roszczenko-Jasińska, Huong N. Vu, Gabriel A. Subuyuj, Ralph Valentine Crisostomo, Elena M. Ayala, James Cai, Nicholas F. Lien, Erik J. Clippard, Richard T. Ngo, Fauna Yarza, Justin P. Wingett, Charumathi Raghuraman, Caitlin A. Hoeber, Norma C. Martinez-Gomez, Elizabeth Skovran

## Abstract

Lanthanide elements have been recently recognized as “new life metals” for diverse environmental microorganisms including Gram-negative methylotrophic bacteria and strains of *Pseudomonas* and *Bradyrhizobium*. Yet much remains unknown regarding lanthanide acquisition and homeostasis. In *Methylorubrum extorquens* AM1, the periplasmic lanthanide-dependent methanol dehydrogenase XoxF1 produces formaldehyde, which is lethal if allowed to accumulate. This property enabled a transposon mutagenesis study to expand knowledge of the metabolic network required for methanol oxidation when lanthanides are available. Growth studies were conducted to detail the involvement of novel gene products that impact the ability of XoxF-type enzymes to oxidize methanol to formaldehyde. The identified genes encode an MxaD homolog, an ABC-type transporter, an aminopeptidase, a putative homospermidine synthase, and two genes of unknown function annotated as *orf6* and *orf7*. Lanthanide transport and trafficking genes were also identified. Growth and lanthanide uptake were measured using strains lacking individual lanthanide transport cluster genes and transmission electron microscopy was used to visualize lanthanide localization. We corroborated previous reports that a TonB-ABC transport system is required for lanthanide incorporation to the cytoplasm. However, cells are able to acclimate overtime and bypass the requirement for the TonB outer membrane transporter to allow expression of *xoxF1* and growth. Transcriptional reporter fusions show that excess lanthanides repress the gene encoding the TonB-receptor. Using growth studies along with energy dispersive X-ray spectroscopy and transmission electron microscopy, we demonstrate that lanthanides are stored as cytoplasmic inclusions that resemble polyphosphate granules.

**IMPORTANCE:** The increasing genetic and biochemical evidence that lanthanide-dependent enzymes are widespread among numerous environmental microbes leads to the parallel questions of how these insoluble metals are scavenged, transported, and used by bacteria. Results herein describe the contribution of the different gene products that constitute the lanthanide utilization and transport machinery in the methylotroph *M. extorquens* AM1 and highlight possible redundancies by periplasmic components. The discovery and characterization of intracellular lanthanide storage in mineral form by these microbes opens the possibility of using methylotrophic platforms for concentration and recovery of these critical energy metals from diverse sources. In addition, methylotrophs are effective biotechnological platforms for the production of biofuels and bioplastics from pollutants such as methane, and inexpensive carbon feedstocks like methanol. Defining the lanthanide acquisition, transport, and storage machinery is a step forward in designing a sustainable platform to recover lanthanides efficiently.

## INTRODUCTION

Lanthanide (Ln) metals have long been recognized for their magnetic and superconductive properties and have facilitated the advancement of our communication, green energy, and medical technologies (1–4). However, it has been less than a decade since an inherent role for Ln in methylotrophic bacteria has been described (5–7). Since these initial reports, bacterial strains that are not considered methylotrophs such as *Pseudomonas putida* and *Bradyrhizobium sp.* have been shown to similarly utilize Ln as cofactors in alcohol dehydrogenase (ADH) enzymes, suggesting the impact of Ln on microbial metabolism may be more widespread than initially thought (8–11). The discovery of Ln-dependent ADH enzymes has led to better identification and characterization of methylotrophs in bacterial communities and the cultivation of new bacterial species in the laboratory (6, 12–14).

Methylotrophic bacteria partake in the global carbon cycle by oxidizing methane, methanol, and other single-carbon substrates, and they have been shown to degrade environmental pollutants like methylmercury (15). They are also industrially important for the conversion of single-carbon waste streams and inexpensive feedstocks into value-added chemicals (16–18). As biological Ln use is a nascent field of study, efforts to understand Ln metabolism will further our understanding of bacterial-mediated environmental processes and community interactions, and can facilitate biological engineering efforts to develop methylotrophic platforms to recover critical Ln metals from waste streams like mining leachate, coal fly ash, and postconsumer electronics (2).

To use methanol as a carbon and energy source, many Gram-negative methylotrophic bacteria first oxidize methanol in the periplasmic space using pyrroloquinoline quinone (PQQ)-dependent ADH enzymes. MxaFI is a two-subunit Ca^2+^-dependent ADH that has been studied for over 60 years and was long considered to be the predominant methanol dehydrogenase (MeDH) in nature until Ln-dependent XoxF enzymes were first described in 2011 (5). When incorporated into the active site, Ln act as potent Lewis acids to facilitate a hydride transfer from the alcohol to the catalytic PQQ cofactor to prompt alcohol oxidation (19–21).

XoxF-type MeDHs were named XoxF before their function was known and based on their homology to MoxF which was the original designation for MxaF (MoxF = MxaF) (6, 19). Since their role in methanol oxidation became apparent, XoxF enzymes have been classified into five phylogenetically distinct clades (Type 1 to 5) (22, 23) yet recent metagenomic sequencing efforts suggest additional XoxF clades may exist (18, 24). In 2016, a Ln-dependent ethanol dehydrogenase was described and named as ExaF based on homology with the Ca^2+^-dependent Exa ethanol dehydrogenases from *Pseudomonas* and *Rhodopseudomonas* strains, and homology with XoxF- and MxaF-type MeDHs (25, 26, 27). While these various ADHs have been designated as MxaFI and XoxF MeDHs and ExaF ethanol dehydrogenases, there is often overlap in the substrate range for the ADHs; XoxF and MxaFI enzymes can oxidize ethanol but have a higher affinity for methanol (19, 22, 23, 25, 28–30), and ExaF ethanol dehydrogenases can oxidize methanol but with less efficiency (26, 27).

All methylotrophic PQQ-ADHs are periplasmic enzymes associated with a cytochrome *c*_L_ (MxaG, XoxG, and ExaG, respectively) that transfers electrons from PQQ to additional cytochromes in the electron transport chain. Operons or genomic clusters that encode *xoxF* and *exaF* genes often contain homologs of *mxaJ* (*xoxJ* and *exaJ* genes respectively), which encode periplasmic binding proteins that have been suggested to function in the activation of the ADHs (31). *mxa* operons encode additional proteins that are suggested to function in Ca^2+^ insertion or to facilitate interactions between the MxaFI MeDH and its cytochrome (MxaACKLD), are required for activation of the *mxa* operon (MxaB), or have unknown function (MxaRSE) (32).

*Methylorubrum extorquens* AM1 (formerly *Methylobacterium extorquens* AM1) produces a MxaFI-type MeDH (encoded by *MexAM1_META1p4538* and *MexAM1_META1p4535*), two XoxF (type 5) MeDHs (encoded by *xoxF1*: *MexAM1_META1p1740* and *xoxF2*: *MexAM1_META1p2757*), and an ExaF-type ethanol dehydrogenase (encoded by *exaF*: *MexAM1_META1p1139*) (26, 27). Before XoxF enzymes were classified into five clades, the XoxF enzymes from *M. extorquens* AM1, which share 90% amino acid similarity, were named as XoxF1 and XoxF2 to distinguish them from one another (33). Phenotypic studies suggest *M. extorquens* AM1 XoxF1 and XoxF2 have redundant function. However, in the laboratory, XoxF1 appears to be the dominant XoxF-MeDH as *xoxF2* is expressed at low levels (25, 27). A fifth putative PQQ-ADH encoded by *MexAM1_META1p4973* does not appear to contribute to methanol growth under laboratory conditions tested (26).

When Ln are absent, MxaFI is the only known contributor to methanol oxidation in *M. extorquens* AM1. When Ln are available, the XoxF enzymes produce formaldehyde from methanol while ExaF oxidizes methanol further to formate (25). Good et al. showed that ExaF can facilitate Ln-dependent methanol growth when *xoxF1* and *xoxF2* are deleted, but at a slower rate (26).

The role of Ln in metabolism is not exclusively catalytic (34–36). When Ln are transported into the cell, a transcriptional response occurs; this effect is referred to as “the Ln-switch” or “rare earth-switch” (27, 37–39). During the Ln-switch, the *mxa* operon (*mxaFJGIRSACKLDEHB*) is downregulated and transcript levels of the *xox1* operon genes (*xoxF1*, *xoxG*, *xoxJ*) are upregulated (25, 27, 37–39).

In *M. extorquens* AM1 and closely related PA1 strain, the MxbDM two-component system along with the *xoxF* genes themselves have been shown to be required for operation of the Ln-switch (33, 40, 41). However, suppressor mutations in the *mxbD* sensor kinase encoding gene can arise, which bypass the need for XoxF1 and XoxF2, presumably by constitutively activating the MxbM response regulator (33, 36, 41). Loss of *xoxF2* alone does not impact growth or operation of the Ln-switch. However, in the absence of *xoxF1*, there is an additive effect when *xoxF2* is deleted from the genome suggesting *xoxF2* may be upregulated to compensate for the absence of *xoxF1* (27, 33). Though much progress has been made regarding the catalytic and regulatory roles of Ln in methylotrophic bacteria, relatively little is known about how these Ln are acquired and incorporated into the enzymes that use them, and if they are stored intracellularly.

The machinery necessary for Ln transport is in the early stages of characterization and is predicted to be analogous to siderophore-mediated iron transport (4, 41) with the siderophore-like molecule referred to as a lanthanophore (42). In *M. extorquens* AM1, ten genes predicted to encode proteins involved in Ln transport and utilization are clustered together in the genome (*MexAM1_META1p1778* – *MexAM1_META1p1787*) and encode an ABC-type transporter, four hypothetical periplasmic proteins of unknown function, a periplasmic protein that binds Ln (encoded by *MexAM1_META1p1781*), a TonB-dependent transporter, lanmodulin (encoded by *MexAM1_META1p1786*), and an additional periplasmic-binding protein capable of Ln binding (4, 43).

Ochsner et al. initially characterized the Ln transport cluster in *M. extorquens* strain PA1 by generating deletions spanning multiple genes in the predicted transport system (41). Their results showed that a TonB-dependent transporter and a putative ABC transporter were necessary for Ln transport. A detailed analysis of the contribution of each gene from the transport cluster is still lacking and Ln transport has not been quantified. Accordingly, Mattocks et al. identified homologs of this system in *M. extorquens* AM1 and showed that a hypothetical periplasmic protein encoded by *MexAM1_META1p1781* can efficiently bind Ln from lanthanum (La^3+^) to gadolinium (4). Additionally, they showed that the periplasmic binding component of the ABC transporter encoded by *MexAM1_META1p1778* was unable to bind Ln, possibly because it instead binds the predicted lanthanophore necessary to transport Ln into the cell.

In this study, we describe new pieces necessary to complete the Ln puzzle: the identification of novel genes that contribute to Ln metabolism and methanol oxidation, and the discovery and visualization of Ln storage in *M. extorquens* AM1. Detailed growth studies for strains lacking each component of the first eight genes in the Ln transport cluster are described, including the ability of these strains to mutate or acclimate (phenotypic change that is inducible and reversible) to allow Ln transport through a predicted secondary mechanism. Ln uptake is also quantified for several transport mutant strains. Finally, we show that *M. extorquens* AM1 stores Ln with phosphate as crystalline cytoplasmic deposits.

## RESULTS

### A genetic study identifies gene products that contribute to methanol oxidation

A transposon mutagenesis study was designed to take advantage of the *in vivo* formaldehyde production capability of XoxF1 and XoxF2 to identify genes required for XoxF-dependent methanol oxidation. The strain used to conduct the mutant hunt contained a mutation in *mxaF* to make cells dependent on the exogenously provided La^3+^ for formaldehyde production, and a second mutation in *fae*, which would result in formaldehyde accumulation and cell death when methanol is oxidized to formaldehyde by XoxF1 and XoxF2 (Fig. 1). Transposon insertions that reduced or eliminated formaldehyde production allowed survival and colony formation on media containing both methanol and succinate, since methanol resistant strains could use succinate for growth. In addition to genes required for XoxF1 and XoxF2 mediated methanol oxidation, this mutant hunt had the potential to isolate insertions in an unknown formaldehyde import system, which would theoretically reduce formaldehyde levels in the cytoplasm (Fig. 1). As ExaF oxidizes methanol directly to formate (25), *exaF* and genes specific to ExaF function were not expected to be identified though this genetic study.

**Fig 1.**
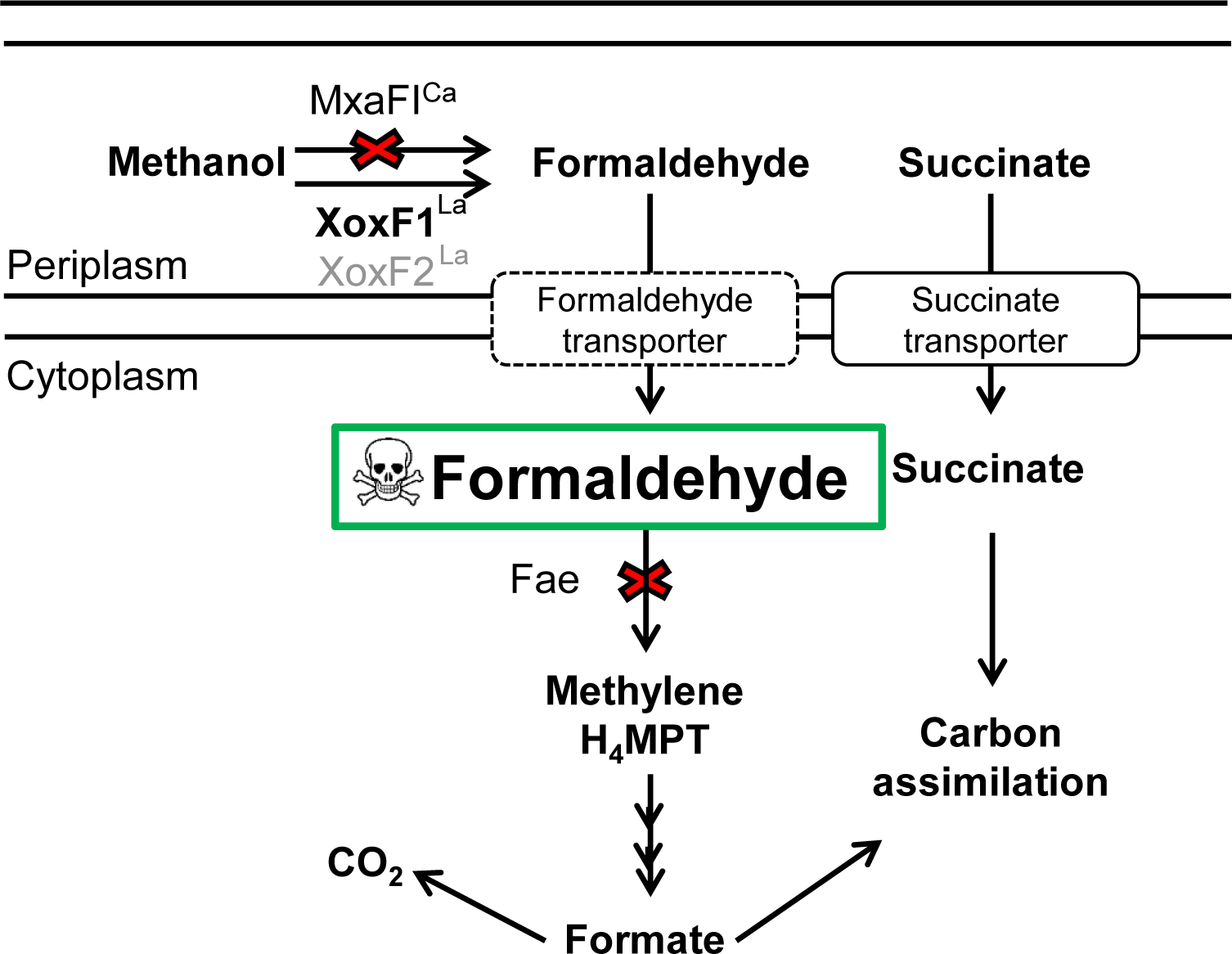
Schematic representation of the metabolic processes relevant to the *mxaF fae* transposon mutagenesis study. XoxF1 and XoxF2 oxidize methanol to formaldehyde, which accumulates to lethal levels in the *fae* mutant strain. If a process required for XoxF-dependent methanol oxidation is disrupted by a transposon insertion, formaldehyde is reduced or eliminated, and cells use succinate for growth. A dashed line around the formaldehyde transporter is to indicate that this function has not been demonstrated.

Over six hundred transposon mutants were isolated, and their insertion locations mapped to the *M. extorquens* AM1 genome. Since it is likely that a portion of these transposon mutants became methanol-resistant due to spontaneous second-site suppressor mutations and not the transposon insertion, only genes that were independently identified four or more times were considered for further analysis and are listed in Table 1. From the twenty eight genes identified in our transposon mutagenesis study (Table 1), mutations in twenty three genes were constructed in *mxaF* and/or wild-type strain backgrounds and methanol growth in the presence of La^3+^ was assessed (Table 2; Table 3).

**Table 1.** Genes identified four or more times via transposon mutagenesis.

| Gene Designation | Gene Name | Predicted Function |
| --- | --- | --- |
| *MexAM1_META1p0863* |  | LysR-type regulator |
| *MexAM1_META1p1292* | *cycL* | *c*-type cytochrome biogenesis |
| *MexAM1_META1p1293* | *cycK* | Heme lyase |
| *MexAM1_META1p1294* | *cycJ* | Periplasmic heme chaperone |
| *MexAM1_META1p1740* | *xoxF1* | Ln-dependent methanol dehydrogenase |
| *MexAM1_META1p1741* | *xoxG* | Cytochrome *c* |
| *MexAM1_META1p1742* | *xoxJ* | Periplasmic binding protein |
| *MexAM1_META1p1746* | *orf6* | Unknown |
| *MexAM1_META1p1747* | *orf7* | Unknown |
| *MexAM1_META1p1748* | *pqqE* | PQQ biosynthesis |
| *MexAM1_META1p1749* | *pqqCD* | PQQ biosynthesis |
| *MexAM1_META1p1750* | *pqqB* | PQQ biosynthesis |
| *MexAM1_META1p1771* |  | MxaD homolog |
| *MexAM1_META1p1778* | *lutA* | ABC transporter-periplasmic binding component |
| *MexAM1_META1p1779* | *lutB* | Exported protein |
| *MexAM1_META1p1782* | *lutE* | ABC transporter-ATP binding component |
| *MexAM1_META1p1783* | *lutF* | ABC transporter-membrane component |
| *MexAM1_META1p1784* | *lutG* | Exported protein |
| *MexAM1_META1p1785* | *lutH* | TonB-dependent receptor |
| *MexAM1_META1p2024* | *hss* | Homospermidine synthase |
| *MexAM1_META1p2330* | *pqqF* | Protease |
| *MexAM1_META1p2331* | *pqqG* | Protease |
| *MexAM1_META1p2359* |  | ABC transporter |
| *MexAM1_META1p2732* | *ccmC* | Heme export |
| *MexAM1_META1p2734* | *ccmG* | *c*-type cytochrome biogenesis |
| *MexAM1_META1p2825* | *ccmB* | Heme export |
| *MexAM1_META1p2826* | *ccmA* | Heme export |
| *MexAM1_META1p3908* |  | Leucyl aminopeptidase |

**Table 2.**
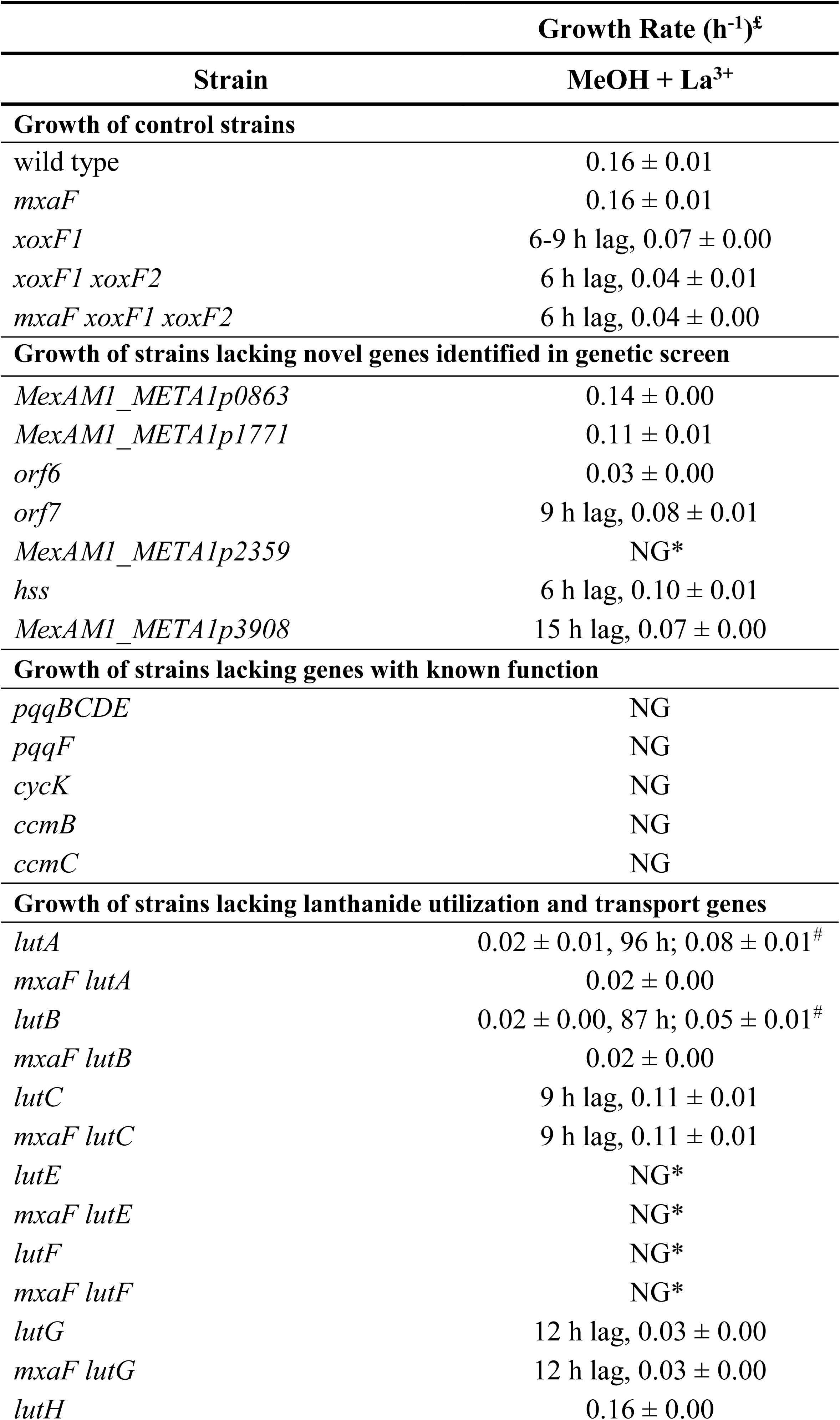
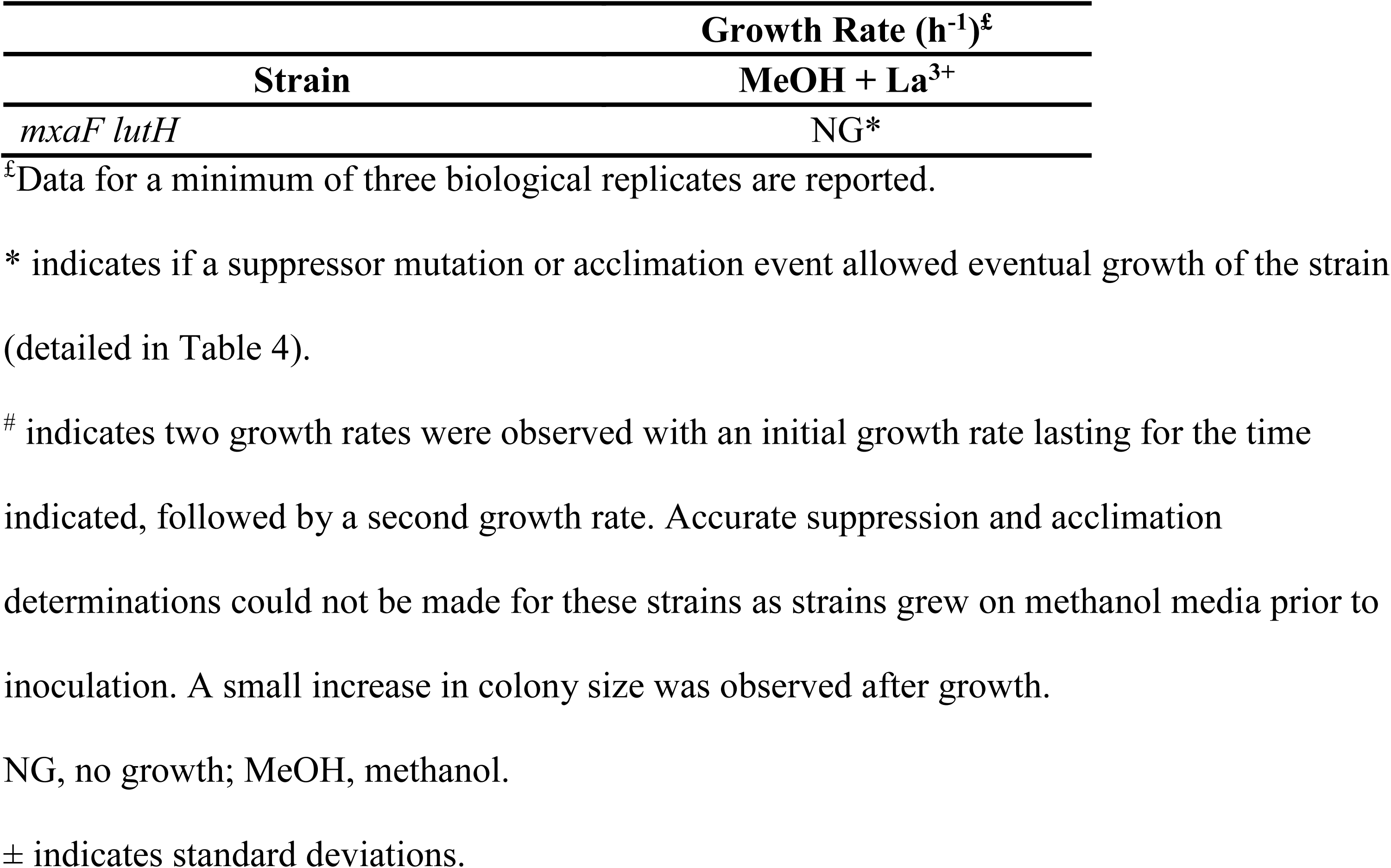
Growth parameters for strains grown in methanol medium with La^3+^.

**Table 3.** Growth parameters for strains grown in methanol medium with and without La^3+^.

| Strain | Growth Rate (h<sup>-1</sup>)<sup>‡</sup> |  |
| --- | --- | --- |
|  | MeOH | MeOH + La<sup>3+</sup> |
| Wild type | 0.14 ± 0.01 | 0.16 ± 0.01 |
| *xoxF1 xoxF2* | NG* | 6 h lag, 0.04 ± 0.01 |
| *xoxG* | 3 h lag, 0.11 ± 0.01 | 0.04 ± 0.00 |
| *xoxJ* | 21 h lag, 0.13 ± 0.00 | 0.04 ± 0.01 |

<sup>‡</sup>Data for a minimum of three biological replicates are reported.
\* indicates a suppressor mutation allowed eventual growth of the strain.
NG, no growth; MeOH, methanol.
± indicates standard deviations.

**Table 4.**
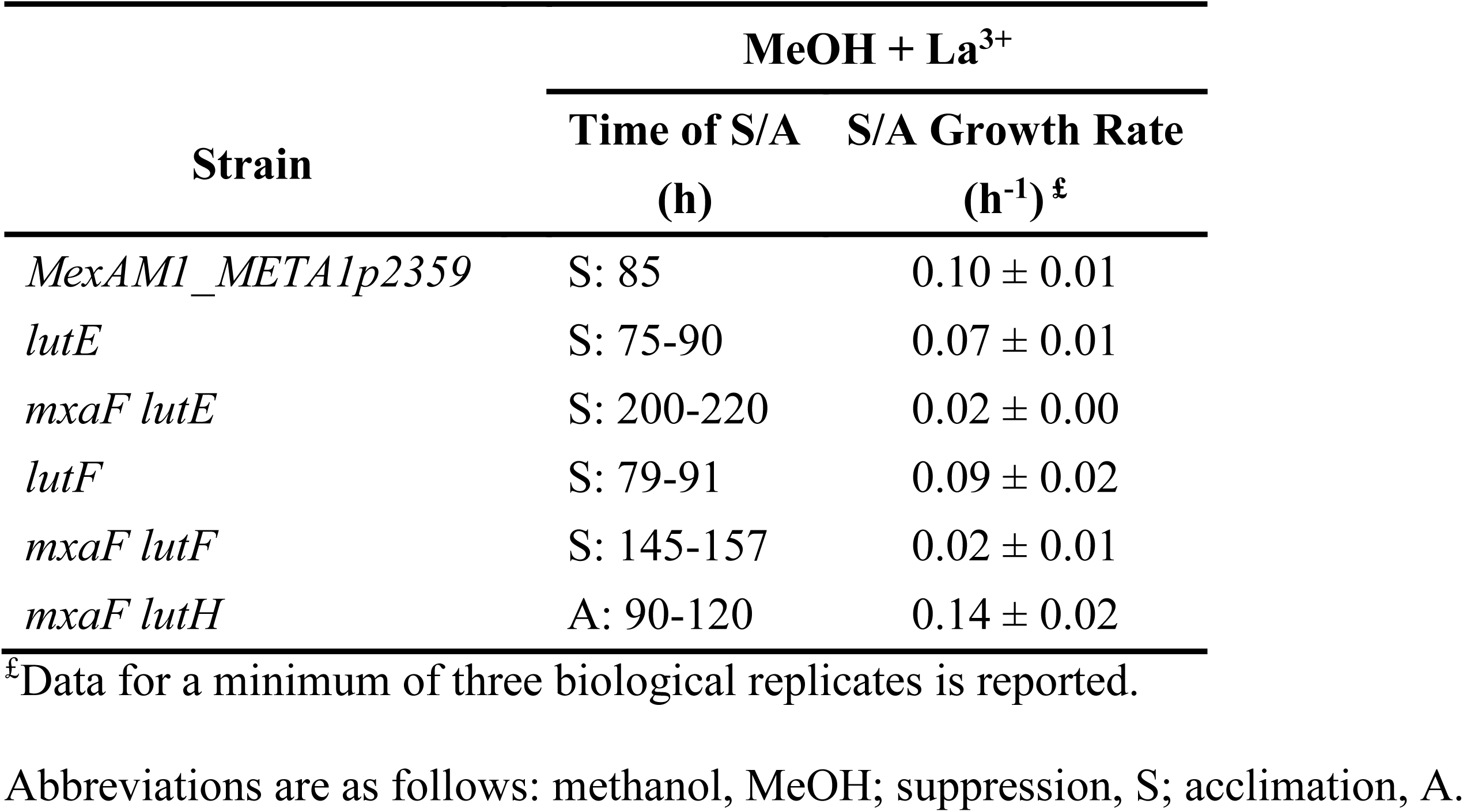
Growth parameters of suppressor and acclimation events.

### Growth phenotypes of strains lacking genes identified in the transposon mutagenesis study

Novel genes which impacted methanol growth when deleted from the *M. extorquens* AM1 genome were identified in the transposon mutagenesis study (Table 1). The identified genes encode a LysR-type transcriptional regulator (*MexAM1_META1p0863*), two proteins of unknown function annotated as *orf6* and *orf7* (*MexAM1_META1p1746* and *MexAM1_META1p1747* respectively), an MxaD homolog (*MexAM1_META1p1771*), a putative homospermidine synthase (*MexAM1_META1p2024*), an ABC-type transporter (*MexAM1_META1p2359*), and an aminopeptidase (*MexAM1_META1p3908*). Growth rates for strains lacking these genes are shown in Table 2 and growth curves for mutant strains that had a 30% or greater reduction in growth rate are shown in Fig. 2. The identification of these genes and initial growth studies lay the foundation to explore the specific roles and requirements for these gene products in XoxF1 enzyme function.

**Fig 2.**
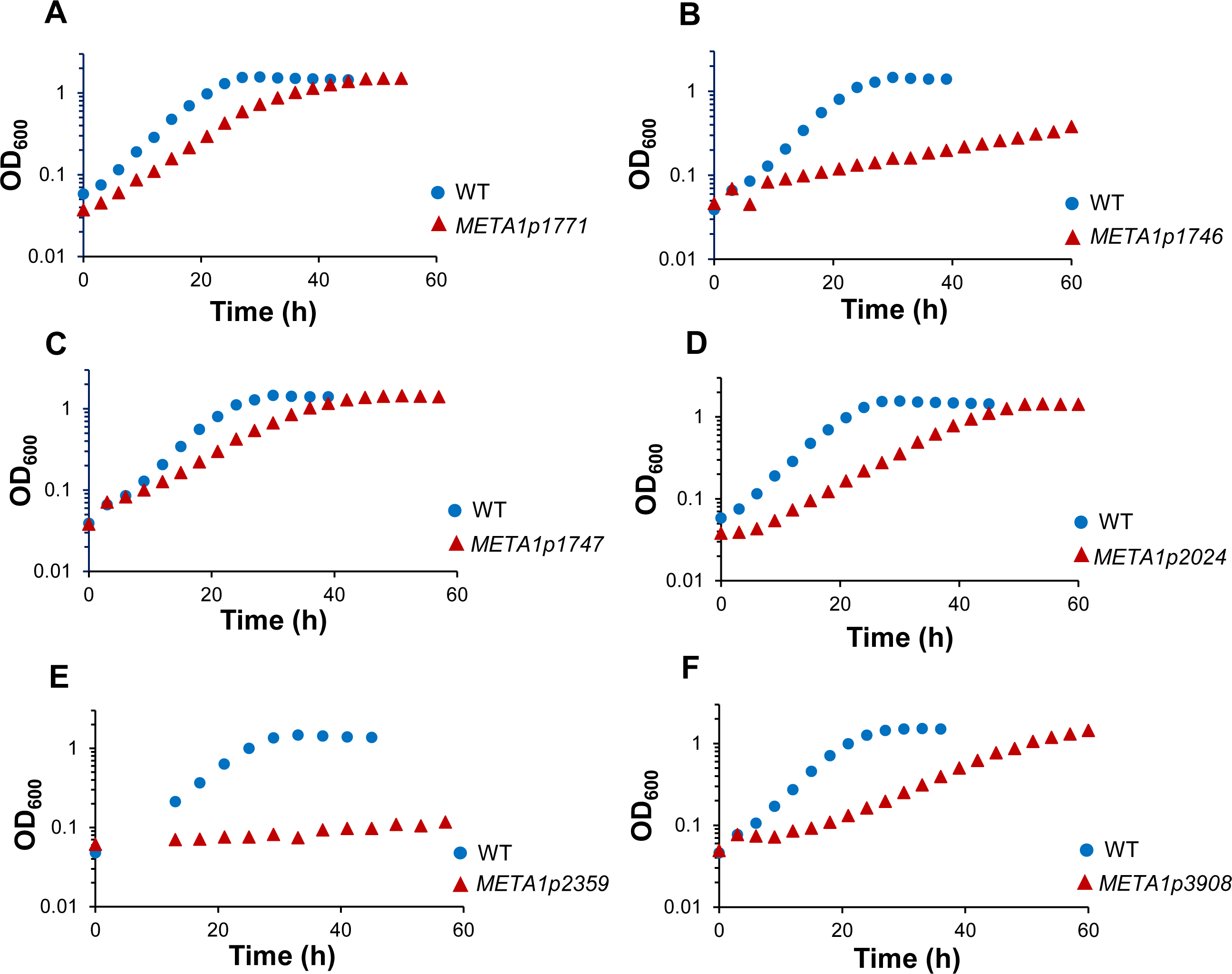
Growth of mutant strains lacking genes identified through transposon mutagenesis in medium containing methanol and La^3+^. Growth of *M. extorquens* AM1 wild-type is represented by blue circles and depicted mutant strains (red triangles) have the following genes deleted: A – *MexAM1_META1p1771*, *mxaD* homolog; B – *MexAM1_META1p1746, orf6*; C – *MexAM1_META1p1747*, *orf7*; D – *MexAM1_META1p2024*, homospermidine synthase; E – *MexAM1_META1p2359*, ABC transporter of unknown function; F – *MexAM1_META1p3908*, aminopeptidase. Representative data from biological triplicates are shown. One-way analysis of variance (ANOVA) determined that growth rate differences between mutant and parent strains are significantly different (*p* < 0.005).

Also included in the list of genes identified in the transposon mutagenesis study were those that encode proteins with described roles in methanol oxidation (PQQ biosynthesis, *xoxF1, xoxG, xoxJ*) and those with predicted functions based on sequence similarity (cytochrome synthesis and Ln transport) (44). Deletion of genes encoding PQQ biosynthesis (Δ*pqqBCDE*; Δ*pqqF*) and each of the three identified cytochrome *c* biogenesis and heme export clusters: *cycK*, heme lyase; *ccmB*, heme exporter; and *ccmC*, heme exporter (reviewed in (44)) eliminated methanol growth as expected (Table 2).

XoxF1 and XoxF2 catalytically contribute to methanol oxidation and growth when methanol and Ln are provided (25, 27), and exert a regulatory role in the absence of Ln as production of XoxF1 or XoxF2 is required for expression of the Ca^2+^-dependent MxaFI-MeDH (27, 33). Growth analysis and transcriptional reporter fusion studies were employed to see if strains lacking *xoxG* or *xoxJ* had similar effects on growth and *mxa* operon expression. In methanol La^3+^ medium, loss of either *xoxG* or *xoxJ* was equivalent to loss of both *xoxF1* and *xoxF2* (Fig. 3; Table 3), which is consistent with XoxG and XoxJ being essential for XoxF-dependent methanol oxidation as suggested by recent biochemical studies (14, 31, 45).

**Fig 3.**
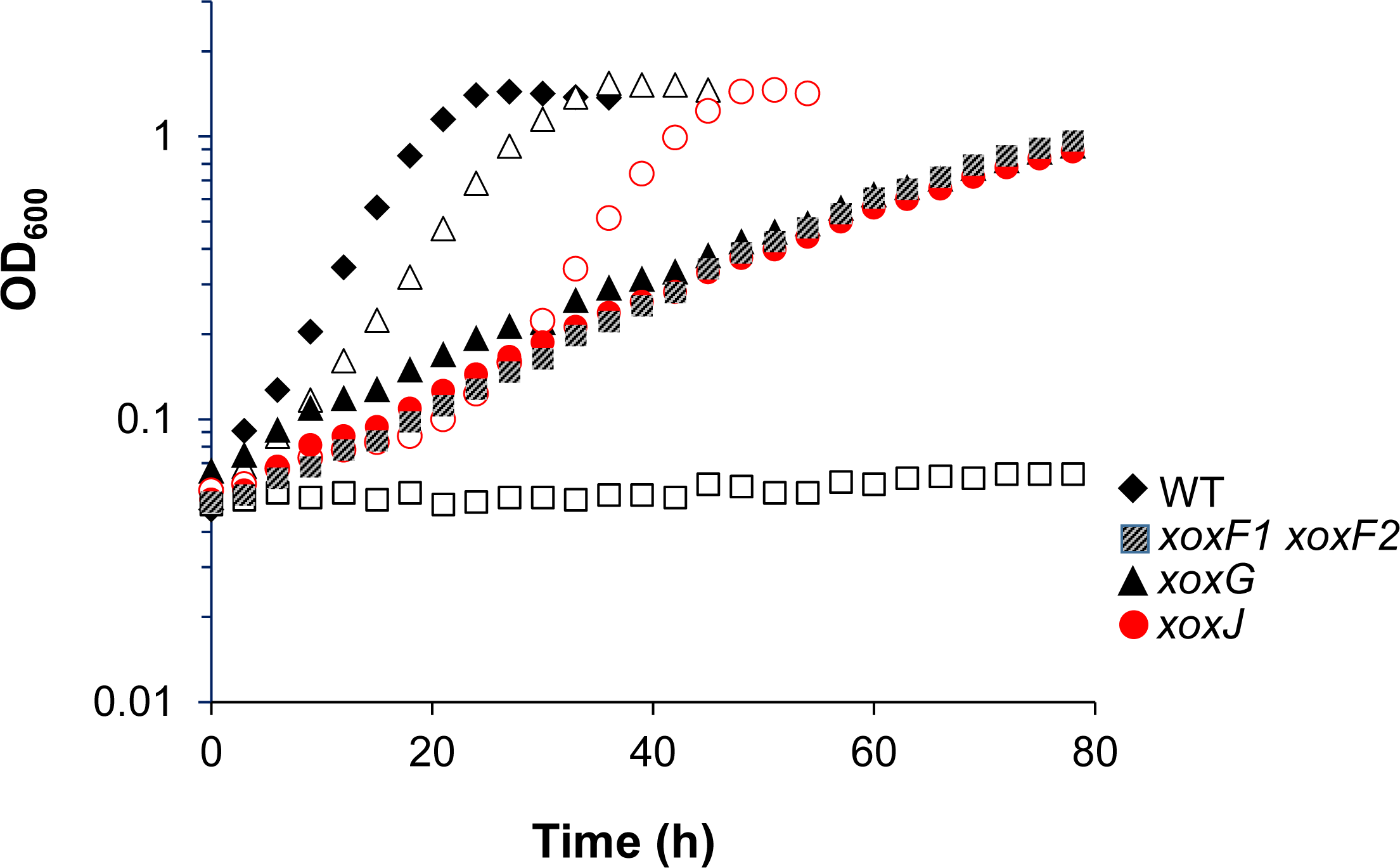
Growth of *xoxG* and *xoxJ* mutant strains in the presence (filled symbols) and absence (open symbols) of La^3+^. Representative data from biological triplicates are shown. One-way analysis of variance (ANOVA) determined that growth rate differences between mutant and parent strains are significantly different (*p* < 0.005).

Unlike the *xoxF1 xoxF2* double mutant strain, transcriptional reporter fusion studies that assessed expression from the *mxa* promoter determined that the growth phenotypes observed for the *xoxG* and *xoxJ* mutants grown in the absence of La^3+^ were not due to impaired *mxa* expression (average relative fluorescence units (RFU)/OD_600_: wild type, 322.9 ± 62.5; *xoxF1 xoxF2*, 3.1 ± 1.3; *xoxG*, 313.9 ± 24.0; *xoxJ*, 512.5 ± 54.9). Taken together, these phenotypes suggest XoxG and XoxJ may play a broader role in methanol metabolism in *M. extorquens* AM1 independent from facilitating XoxF1 and XoxF2 catalytic and regulatory functions.

### Detailed phenotypic characterization of Ln utilization and transport cluster mutant strains

Consistent with previous reports for strain PA1, our transposon mutagenesis studies suggest that genes encoding homologs of the TonB- and ABC-dependent Fe^3+^ scavenging systems play a role in methanol metabolism when Ln are present (41). In addition to the transport system homologs, two of six hypothetical periplasmic proteins encoded in the Ln transport cluster were identified in the transposon mutagenesis study (Fig. 4A). We expanded Ochsner’s study by dissecting the contribution of individual genes (*MexAM1_META1p1778* through *MexAM1_META1p1785*). Further, our results show that La^3+^ transport can occur in the absence of the TonB-ABC transport system by acclimation and/or suppression. Based upon work detailed within, we propose to name the Ln transport cluster genes as *lut*, for **<u>L</u>**n **<u>u</u>**tilization and **t**ransport.

**Fig 4.**
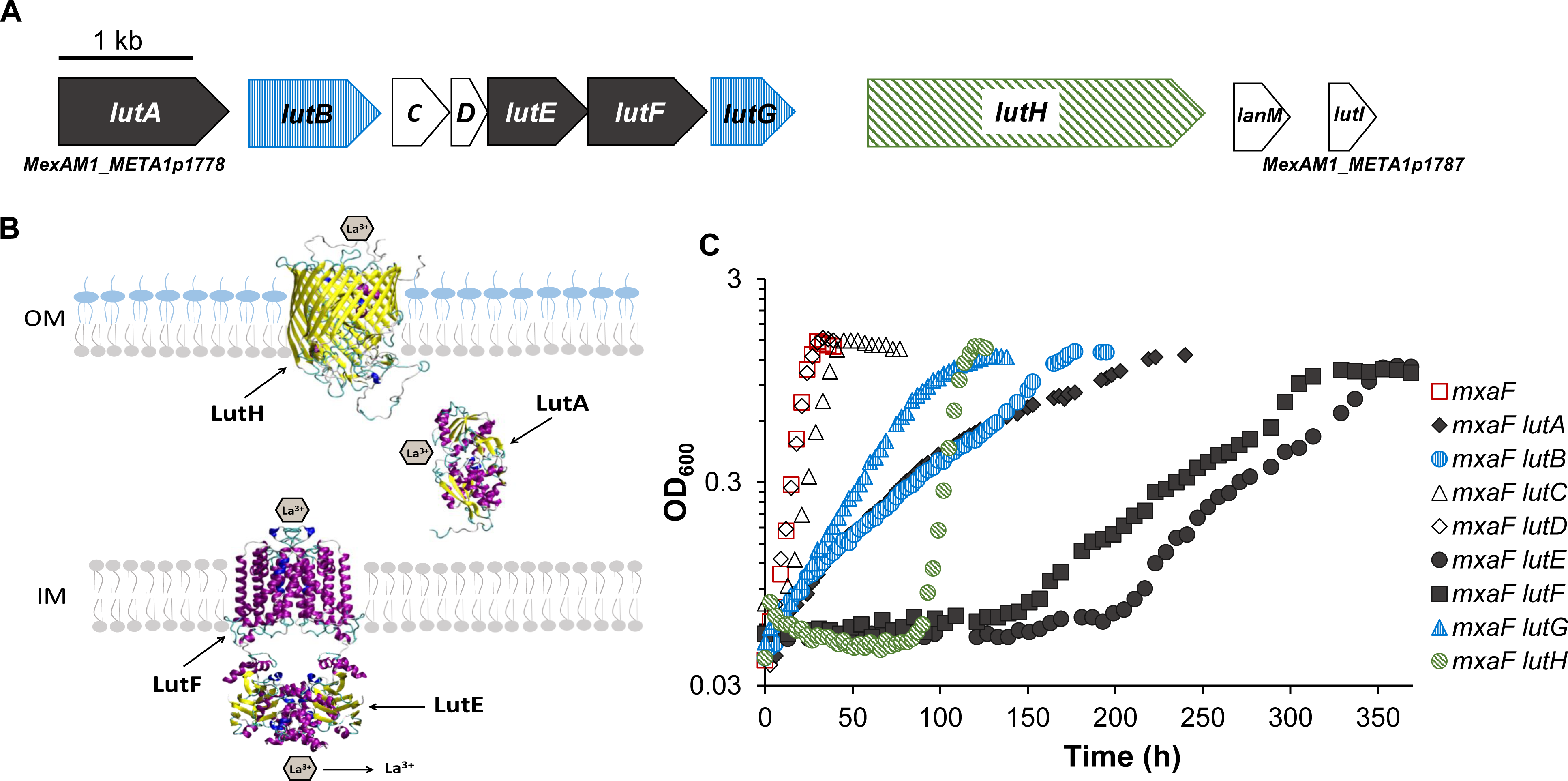
Characterization of the *lut* cluster. A) Genomic map of the *lut* genes (*MexAM1_META1p1778* to *MexAM1_META1p1787*). Black, genes encoding the ABC transport system; green, gene encoding the TonB-dependent transporter; blue, genes encoding putative periplasmic proteins identified by transposon mutagenesis; white, lanmodulin and additional genes encoding periplasmic proteins not identified by transposon mutagenesis. B) Model for Ln transport. The three-dimensional structures of monomers of LutH and LutA were predicted using homology modeling (HHpredserver and MODELLER) (89) and homodimers of the ABC transporter were predicted using GalaxyHomomer (90). C) Growth of *mxaF* and *mxaF lut* mutant strains with 2 μM LaCl_3_. Graphs depict representative data from three biological replicates. Growth rate averages and standard deviations are shown in Table 2.

As reported by Ochsner et al. (41), loss of *lutH* (*Mextp1853* in strain PA1) alone did not result in a growth defect in medium containing methanol and La^3+^, which is consistent with the hypothesis that the TonB-dependent transporter is needed for transport of La^3+^ into the periplasm. If La^3+^ does not enter the periplasm, the *mxaF* and *mxaI* genes are likely expressed and used for methanol oxidation. Loss of both *mxaF* and *lutH* arrested growth, but after 90-120 h, growth of the *mxaF lutH* double mutant strain occurred in approximately 60% of the 24 cultures tested (Fig. 4C; Table 2). Consistent with an acclimation process and not a suppressor mutation, the *mxaF lutH* double mutant strain lost the ability to grow in methanol medium with La^3+^ after passage onto medium containing succinate. To determine if acclimated growth was due to production of XoxF1, expression from the *mxa* and *xox1* promoters was measured before and after acclimation. Before acclimation, *mxa* promoter expression occurred (RFU/OD_600_: 363 ± 11) while *xox1* expression was repressed as the cells could not transport La^3+^ to promote the Ln-switch. Once acclimation occurred and the strain began to grow, expression from the *mxa* promoter was repressed and expression from the *xox1* promoter increased dramatically from RFU/OD_600_: 53 ± 7 to 1140 ± 73, suggesting the cells were able to uptake La^3+^ through the outer membrane via an unknown mechanism.

LutA, a periplasmic binding protein, is predicted to traffic lanthanophore bound Ln through the periplasm to the inner membrane transporter system (encoded by *lutE* and *lutF*) for transport into the cytoplasm (Fig. 4B) (43). Strains lacking *lutE* or *lutF* in the *mxaF* mutant strain background were unable to grow in medium containing methanol and La^3+^ (Fig. 4C). However, after 200 h and 150 h respectively, second-site suppressor mutations arose which facilitated growth albeit 88% slower than that of the wild-type strain (Fig. 4C; Table 2). In contrast, loss of *lutA* in the absence of *mxaF* still allowed La^3+^-dependent growth but at a reduced rate. Genes *lutB* and *lutG*, encoding hypothetical periplasmic proteins, were also identified in the transposon mutagenesis study. Growth of the *mxaF lutB* and *mxaF lutG* double mutant strains was similar to that of the *mxaF lutA* double mutant strain, suggesting an equally important but non-essential function. The growth observed for the *lutA*, *lutB*, and *lutG* mutant strains in the absence of *mxaF* was not identified as acclimation or second-site suppression. To ensure that observed phenotypes were not due to polarity, mutants lacking individual transport cluster genes (*lutABEFG*) were complemented by expressing the respective gene in pCM62 (46) and growth similar to the wild-type strain was restored in each case (Fig. S1).

Genes encoding LutC, LutD, and LanM were not identified in the transposon study but LutD and LanM have been shown to bind Ln (4). Loss of *lutD* did not result in a significant growth defect in the *mxaF* mutant and wild-type strain backgrounds while loss of *lutC* resulted in a small growth defect when La^3+^ was provided (Fig. 4C; Table 2).

### Quantification of La^3+^ uptake

To assess how loss of the Ln transport system affects La^3+^ uptake, La^3+^ levels were quantified from the spent media at different phases of growth (Fig. 5). Strains were grown in methanol medium containing 2 µM LaCl_3_ and limiting succinate (3.75 mM) as *lutA, lutE*, and *lutF* mutant strains are unable to grow or grow poorly with methanol as a sole carbon source (Fig. 4C; Table 3). While significant decreases in La^3+^ levels were not observed for any strain during early- to mid-exponential phase (Fig. 5B), as cultures continued to grow, and succinate was likely depleted, significant differences in La^3+^ levels became apparent. In the wild-type strain, La^3+^ uptake continued into stationary phase (Fig. 5B). Growth of the *lutE* and *lutF* mutants (encoding ABC-transporter components) was arrested at an approximate OD_600_ of 0.7, yet La^3+^ concentrations in the spent media continued to decrease to 0.7 µM eight hours into stationary phase (Fig. 5A). Taken together with the growth phenotypes, these data suggest that in the absence of the Lut-ABC-transporter system, Ln are likely transported through the outer membrane but cannot enter the cytoplasm. Levels of La^3+^ in the supernatants from the *lutH* mutant (*lutH* encodes the TonB-dependent transporter) were significantly higher than wild type in mid-exponential and stationary phases (Fig. 5B). A decrease in La^3+^ content from the *lutH* mutant strain culture supernatants was unexpected but may reflect adsorption of La^3+^ onto the surface of the cells or interaction of La^3+^ with lipopolysaccharide, a phenomenon observed for other metals in different bacterial species (47).

**Fig 5.**
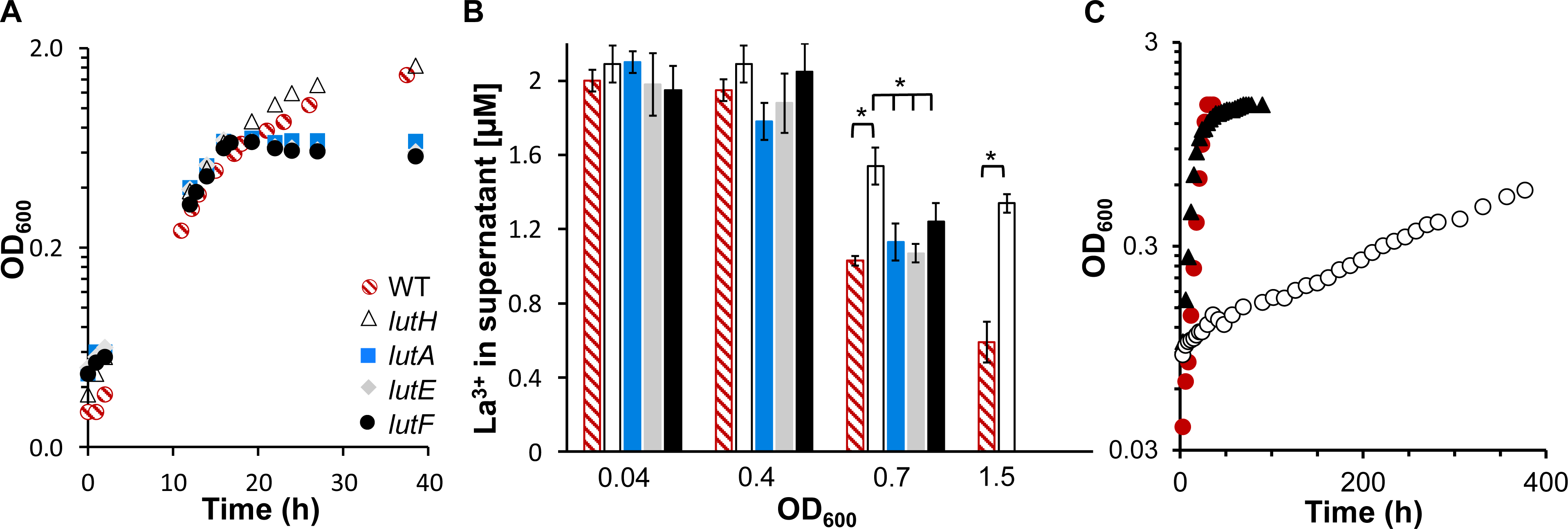
Growth studies and measurement of La^3+^ uptake suggest La^3+^ storage capability. A) Growth of wild type (red dashed circles), *lutH* (white triangles), *lutA* (blue squares), *lutE* (gray diamonds), and *lutF* (black circles) strains grown with limiting succinate (3.75 mM), methanol (125 mM), and 2 μM LaCl_3_. Data are the average of three biological replicates. Standard deviations between biological replicates were less than ±0.02. B) La^3+^ concentrations (μM) in culture supernatants of *lut* mutant strains depicted in same color code as in (A). Data depict the average of three biological replicates with error bars showing the standard deviation. One-way analysis of variance (ANOVA) followed by a T-test was used to represent statistical significance. *The *p*-value is <0.005. C) Growth of the *mxaF* mutant strain with excess LaCl_3_ (20 mM) (red filled circles). Stationary phase cells were washed and reinoculated into media lacking Ln (subculture #1, black triangles) and the process was repeated (subculture #2, empty circles). Representative data from three biological replicates is shown. Standard deviations for growth rates between replicates were less than ± 0.02.

### Expression from the *lutH* promoter is repressed by La^3+^

Genes found within the *lut* cluster are likely not expressed as a single transcript as suggested by the spacing between genes like *lutG* and *lutH*, and by multiple promoters predicted by the bacterial promoter prediction tool, BPROM (48). To test if expression of the of Ln uptake genes is regulated in a similar manner to Fe^3+^ uptake genes, a fluorescent transcriptional reporter fusion was used to monitor expression from the predicted *lutH* promoter region in methanol media. Addition of exogenous La^3+^ repressed expression 4-fold (RFU/OD_600_: 210 ± 7 without La^3+^ and 47 ± 5 with La^3+^) showing that when Ln are in excess, transport is down-regulated. This is consistent with the mechanism of control for iron homeostasis (49, 50). Further, during the revision of this manuscript, similar repression was reported for transcription of the TonB transporter of *Methylotuvimicrobium buryatense* 5GB1C (51).

### La^3+^ is stored as cytoplasmic crystalline deposits

The continued uptake of La^3+^ in stationary phase suggested the possibility of Ln storage. To determine if La^3+^ storage could facilitate methanol growth in the absence of exogenously supplied Ln, an *mxaF* mutant strain was grown in methanol medium with 10-times the standard concentration of LaCl_3_, resulting in a growth rate of 0.15 ± 0.00 h^−1^. Once in stationary phase, cells were washed and sub-cultured into methanol medium lacking La^3+^. As shown in Fig. 5C, a similar growth rate to that observed when cells were grown with La^3+^ was obtained until the culture reached an approximate OD_600_ of 1.0. A slower growth rate was noted as growth continued to an OD_600_ of 1.5. After a second subculture into media lacking La^3+^, cells continued to grow but at a severely impaired rate (0.01 ± 0.0 h^−1^) until they reached a peak OD_600_ of 1.3. These data are consistent with the ability of Ln-storage to supply Ln for methanol oxidation until the storage is depleted. As these storage growth curves were carried out for >400 h, a control experiment was done to determine growth due to either background levels of non-MxaF ADH activity or due to leaching of residual Ln from the glassware. A growth rate of 0.004 ± 0.001 h^−1^ was observed when an *mxaF* mutant strain that had not been previously grown with La^3+^ was tested.

Transmission electron microscopy (TEM) coupled with energy dispersive X-ray spectroscopy (EDS) has been used to determine the elemental composition of cellular inclusions (52, 53) while La^3+^ has been widely used as an intracellular and periplasmic stain for electron microscopy (54–56). Here, we show that La^3+^ can be directly identified by TEM if accumulated inside *M. extorquens* AM1 cells. Electron-dense deposits were observed in the cytoplasm from *M. extorquens* AM1 cells grown with exogenous LaCl_3_ (Fig. 6A-B). Samples were analyzed using EDS and corroborated that the electron dense deposits contained La^3+^ (Fig. 6C). When grown without La^3+^, only a few cells showed smaller electron dense areas (Fig. 6E-F); however, La^3+^ was not detected in EDS analysis in these cases (Fig. 6G). These data demonstrate that La^3+^ can be stored in *M. extorquens* as metal deposits. Moreover, EDS analysis of electron dense areas from the wild-type strain grown with La^3+^ determined a content of lanthanum (22.2 ± 1.0%), phosphorous (15.1 ± 2.1%), and oxygen (51.1 ± 1.9%) suggesting La^3+^ is complexed with phosphate (Fig. S2). Traces of chloride (3.0%), calcium (2.2%), and aluminum (3.4%) ions were also detected. The copper, carbon, and silicon ion content from the support grids and embedding medium were not considered for metal content calculations. High-resolution transmission electron microscopy (HRTEM) images of the La^3+^ deposits showed an atomic lattice with a Moiré fringes pattern, indicating a crystalline nature (57) (Fig. 6D). Together, these results suggest that La^3+^ is embedded in inorganic phosphate crystals, which form the electron dense deposits observed in the cytoplasm.

**Fig 6.**
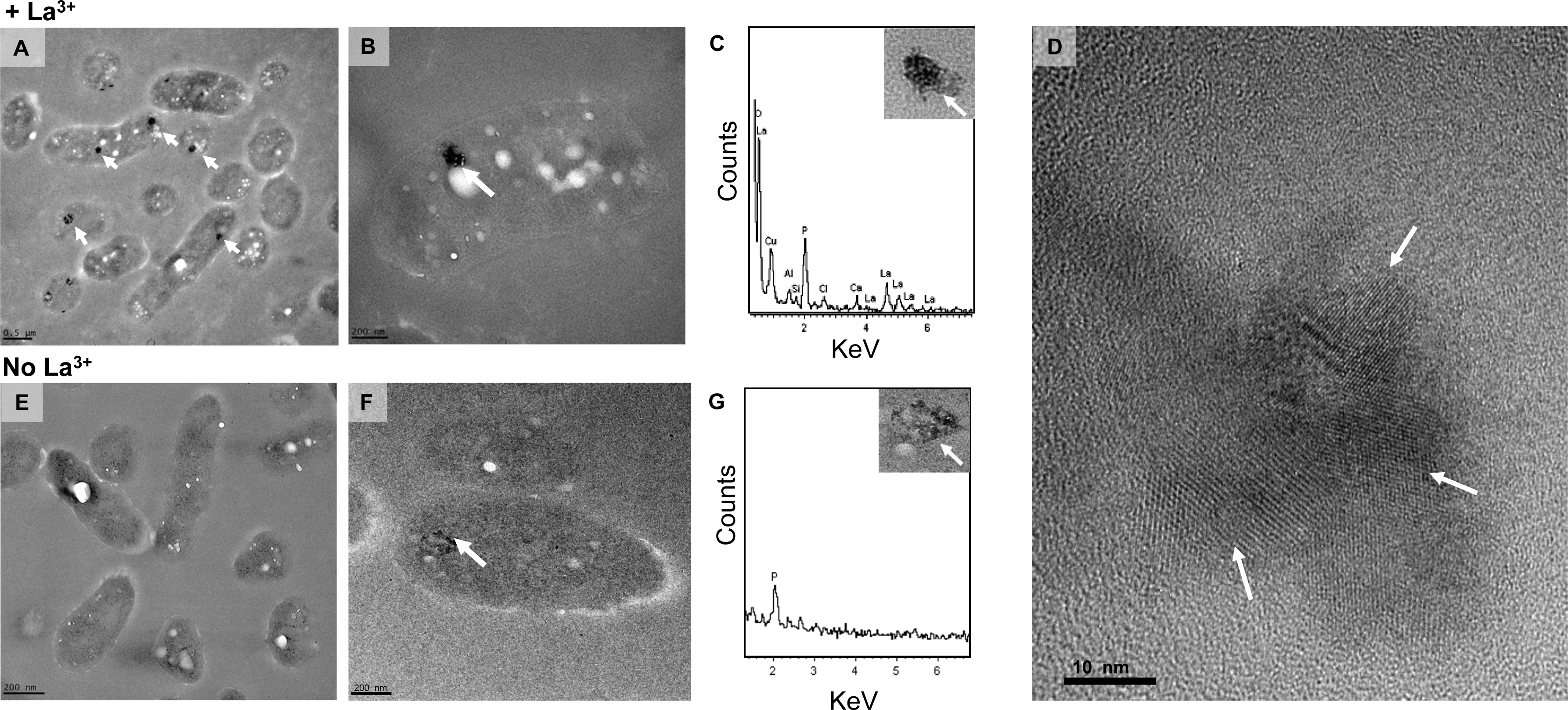
Visualization of La^3+^ storage. TEM of ultrathin sections of wild-type cells grown with 3.75 mM succinate and 125 mM methanol containing (A-B) and lacking (E-F) 20 µM La^3+^. C) and G) Elemental analysis of electron dense deposits from cells grown with (C) and without (G) La^3+^. D) High-resolution transmission electron microscopy analysis of the wild-type strain shows an atomic lattice structure in electron-dense areas suggesting that La^3+^ is embedded in cytoplasmic crystals.

### Visualization of La^3+^ accumulation in *lut* transporter mutants

To determine if La^3+^ could be visualized in strains lacking the Lut-TonB-ABC transport system, TEM and EDS were employed for the analysis of *lutA*, *lutE*, *lutF*, and *lutH* mutant strains. Strains were grown in methanol medium containing La^3+^ and limiting succinate. Samples stained with OsO_4_ and 2% uranyl acetate allowed the outer and inner membrane of the bacterial cells to be distinguished (Fig. 7A-E left subpanels), while visualization without staining enabled metal content analysis by removing the interaction between Os^8+^ and phosphate, which interferes with La^3+^ measurements (Fig. 7A-E right subpanels). Mutants lacking the TonB-dependent transporter (encoded by *lutH*) did not display La^3+^ deposits (Fig. 7A). In contrast, localized La^3+^ deposits were visualized in the periplasmic space in mutant strains lacking the ABC-transporter components (encoded by *lutA, lutE*, and *lutF*) as shown in Fig. 7B-E. EDS microanalyses confirmed these electron dense periplasmic deposition areas contained La^3+^ (Fig. 7F). Taken together, these findings directly demonstrate a role for the TonB-dependent and ABC transporters in Ln transport.

**Fig 7.**
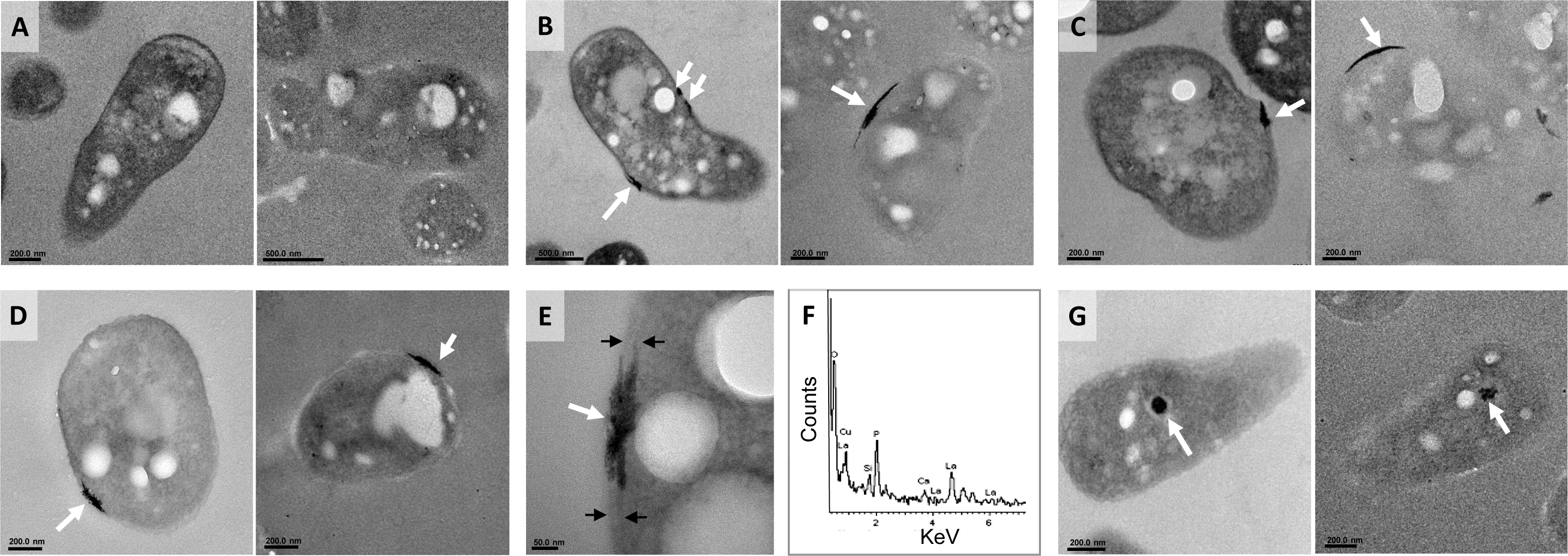
Visualization of La^3+^ localization using TEM. Thin sections of A) *lutH*, B) *lutE*, C) *lutF*, D) *lutA*, and G) wild-type strains grown with 3.75 mM succinate, 125 mM methanol, and 20 µM LaCl_3_. White arrows indicate deposits of electron scattering material in the periplasm. Cells were fixed with 2.5% glutaraldehyde and stained with OsO_4_ and uranyl acetate to detect cell membranes (left subpanel), or left unstained for elemental analysis (right subpanel). E) Magnification of the La^3+^-deposits localized in the periplasmic space from the *lutA* mutant strain; black arrows indicate the boundaries of the outer membrane and inner membrane. F) Elemental analysis of the electron-dense deposits observed inside the periplasm.

## DISCUSSION

The Ln-dependent XoxF1-MeDH has been shown to produce formaldehyde *in vivo* (25). Here, we took advantage of this property which allowed lethal levels of formaldehyde to accumulate when methanol was oxidized by the XoxF enzymes and *fae* was deleted from the genome. These phenotypes enabled a genetic selection to identify gene products required for or involved in methanol oxidation by the XoxF enzymes.

Novel genes were identified that when deleted from the genome, resulted in a 30% or greater reduction in growth rate in medium containing methanol and La^3+^. An ABC-transporter of unknown function (encoded by *MexAM1_META1p2359*) was found to be essential for methanol growth. Some possible functions for META1p2359 may include formaldehyde import, or export of Ln or PQQ into the periplasm for incorporation into the ADHs. Further, a gene encoding a putative homospermidine synthase (*hss*) was identified. Homospermidine synthases function in polyamine biosynthesis and their products (e.g. putrescine, spermidine) have been implicated in roles as diverse as host pathogen interactions, biofilm formation, siderophore production, acid resistance, and free radical scavenging (58, 59).

Before the existence of Ln-dependent MeDHs was known, it was concluded that *orf6* and *orf7* gene products did not have a role in C_1_-metabolism though *orf6* and *orf7* were proximal to other methylotrophic genes (60). Results here clearly suggest that *orf6* and *orf7* are important for Ln-dependent methylotrophy.

A homolog of MxaD (encoded by *MexAM1_META1p1771*) was also shown to be important for methanol La^3+^ growth. MxaD is a 17-kDa periplasmic protein that directly or indirectly stimulates the interaction between the MxaFI-MeDH and cytochrome *c*_L_ (61). Interestingly, upstream of *MexAM1_META1p1771* is another *mxaD* homolog (*MexAM1_META1p1772*) which was not identified in the transposon mutagenesis study. The presence of multiple *mxaD* homologs may suggest redundancy or that the various MxaD homologs may function with different ADH enzymes.

As expected, transposon insertions were found within genes with known or predicted roles in methanol oxidation like PQQ biosynthesis, cytochrome biogenesis, and the *xoxF_1_GJ* genes. Recent biochemical and structural analyses of XoxG suggest that this cytochrome is tuned specifically for light Ln (lanthanum to samarium), while XoxJ interacts with and may activate XoxF (31). Here we show that loss of either *xoxG* or *xoxJ* resulted in growth that mirrored the *xoxF1 xoxF2* double mutant strain, consistent with XoxG and XoxJ as essential for the activity of XoxF-enzymes.

Using growth and transcriptional reporter fusion studies, we show that unlike XoxF1, XoxG and XoxJ are not required for expression of the *mxa* genes yet loss of the *xoxG* and *xoxJ* genes impacts growth in the absence of La^3+^. In the methanotroph *Methylomonas* sp. strain LW13, loss of *xoxG* also results in a growth defect in methanol medium lacking Ln suggesting an unknown role in metabolism in addition to functioning as a cytochrome for XoxF-mediated methanol oxidation (45). Our results are in contrast to previous reports for *M. extorquens* PA1 and AM1, where loss of *xoxG* and *xoxJ* did not result in a growth phenotype in methanol medium lacking Ln (41, 60). Notably, the previous AM1 studies were carried out on agar plates where subtle growth defects may not be apparent, and the growth media used for the previous PA1 and AM1 studies was not the same as used in these studies.

Growth phenotypes for *M. extorquens* AM1 PQQ biosynthesis mutants have been reported previously in the absence of Ln (62) but *ccm* and *cyc* mutants have not been characterized and the functions encoded by these genes inferred from sequence homology. Our results using La^3+^ corroborate that the biosynthesis of PQQ, heme export, and cytochrome biogenesis enable Ln-dependent ADH activities as predicted.

Many insertions identified in this study mapped to the previously identified Ln transport gene cluster (4, 41). Characterization of strains lacking genes in the *lut* cluster is consistent with the observations recently reported for *M. extorquens* PA1 (41) where a TonB-ABC transport system contributes to Ln transport into the cytoplasm. It has been proposed that Ln may be acquired via lanthanophores (42), which enter the periplasm through the TonB-dependent transporter (4, 41). However, the acclimation of the *mxaF lutH* double mutant strain allowing *xoxF1* expression and growth suggests that an additional mechanism exists to take up Ln. Additionally, we show that similar to the paradigm for regulation of siderophore mediated Fe^3+^ uptake, expression from the TonB-dependent transporter promoter, *lutH*, is repressed by La^3+^.

Since transposon mutants disrupted in the genes encoding lanmodulin or LutD were not isolated, our observations are consistent with the findings in *M. extorquens* PA1 that suggest a non-essential or redundant role for these proteins in Ln metabolism (41). Notably, the *mxaF fae* transposon mutagenesis study did not identify obvious gene candidates for lanthanophore biosynthesis though over 600 insertions were mapped to the genome. This may indicate that more than one lanthanophore is produced by the cell or that the lanthanophore has an essential role that is not yet understood. Additionally, mutations in the TonB-ExbB-ExbD energy transducing system genes were not identified. Homologs are encoded in the *M. extorquens* AM1 genome in single copy (*MexAM1_META1p1510 – MexAM1_META1p1512*). Using a variety of media conditions, repeated attempts were made to delete TonB without success, suggesting TonB may be essential for *M. extorquens* AM1.

Our transport, TEM, and EDS analyses are consistent with LutH facilitating Ln transport into the periplasm and the Ln-ABC transport system facilitating Ln transport into the cytoplasm. La^3+^ concentrations found in the supernatant from strain variants lacking transport system components suggest that once in the periplasm, significant concentrations of La^3+^ do not go back outside of the cell as ABC transporter mutant strains showed uptake of La^3+^ from the medium similar to the wild-type strain. Intriguingly, TEM and EDS studies with ABC transporter mutant strains demonstrated localized accumulation of La^3+^ in the periplasmic space. It is not yet clear how or why La^3+^ accumulate in specific areas rather than appear diffused throughout the periplasm. Taken together, our phenotypic growth, transport, and visualization studies suggest that Ln must enter the cytoplasm for the *xoxF* and *exaF* ADHs to be expressed and that Ln uptake into the periplasmic space do not support methanol dehydrogenase activity but may be enough to repress *mxa* expression. This work will facilitate new exploration of the Ln-switch.

It has been observed that bacteria such as *Bacillus licheniformis* (63), *Myxococcus xanthus* (56), and *Pseudomonas aeruginosa* (64) can effectively adsorb Ln in mineral form onto their cell surface when Ln are at high concentrations (mM). Unlike these microorganisms, our TEM and EDS analyses demonstrate that *M. extorquens* AM1 stores Ln in the cytoplasm in crystal form. This is consistent with recent studies for the non-methylotroph *Thermus scotoductus* SA-01 which showed that europium (Eu^3+^) can accumulate in the cytoplasm. However, these studies were conducted using very high Eu^3+^ concentrations that are not typically found in nature (53). Further, a role for Eu in the metabolism of *T. scotoductus* SA-01 remains to be defined.

For many bacteria, biomineralization is a mechanism used to cope with toxicity of different metals, manage waste products, sense and change orientations in accordance with geomagnetic fields, and store important cations for growth (65–70). It has been reported that some microorganisms store cations like Mg^2+^ and Ca^2+^ complexed to polyphosphate in the form of volutin (also known as metachromatic granules) or acidocalcisomes (71–74). It is not yet known if *M. extorquens* AM1 stores La^3+^ complexed to polyphosphate, however, the ratios of P, O, and La^3+^ detected in our studies are consistent with La^3+^ phosphates. Detailed studies are necessary to define the exact chemical structure of Ln storage deposits in *M. extorquens* AM1 and if these granules are membrane or lipid bound. Our current findings bring exciting implications for bacterial metabolism and cell biology, and for the development of bioremediation and biometallurgy strategies for Ln recovery.

## MATERIALS AND METHODS

### Bacterial strains and cultivation

Strains and plasmids used in this study are listed in Table 5. *Escherichia coli* strains were cultivated in Lysogeny Broth (LB) medium (75) (BD, Franklin Lakes, NJ) at 37°C. *M. extorquens* AM1 strains were grown in *Methylobacterium* PIPES [piperazine-*N,N*’-bis(2-ethanesulfonic acid)] (MP) media (76) supplemented with succinate (15 mM) and/or methanol (125 mM) as described (26) unless otherwise stated. Conjugations took place on Difco Nutrient Agar (Thermo Fisher Scientific, Waltham, MA). Cultures were grown in round-bottom polypropylene or borosilicate glass culture tubes, or 250 mL polypropylene Erlenmeyer flasks (Thermo Fisher Scientific, Waltham, MA). If glass tubes or flasks were used to culture bacteria, they were pretreated to remove Ln as previously described (27). Liquid cultures were grown at 29°C and shaken at 200 and 180 rpm in New Brunswick Innova 2300 and Excella E25 shaking incubators (Eppendorf, Hauppauge, NY), respectively. LaCl_3_ was supplemented to a final concentration of 2 or 20 μM when indicated. When necessary, antibiotics were added at the following concentrations: rifamycin (Rif, 50 µg/mL), tetracycline (Tc, 10 µg/mL for LB, 5 µg/mL for MP or 10 µg/mL when used together with Rif), kanamycin (Km, 50 µg/mL), ampicillin (Ap, 50 µg/mL).

**Table 5.**
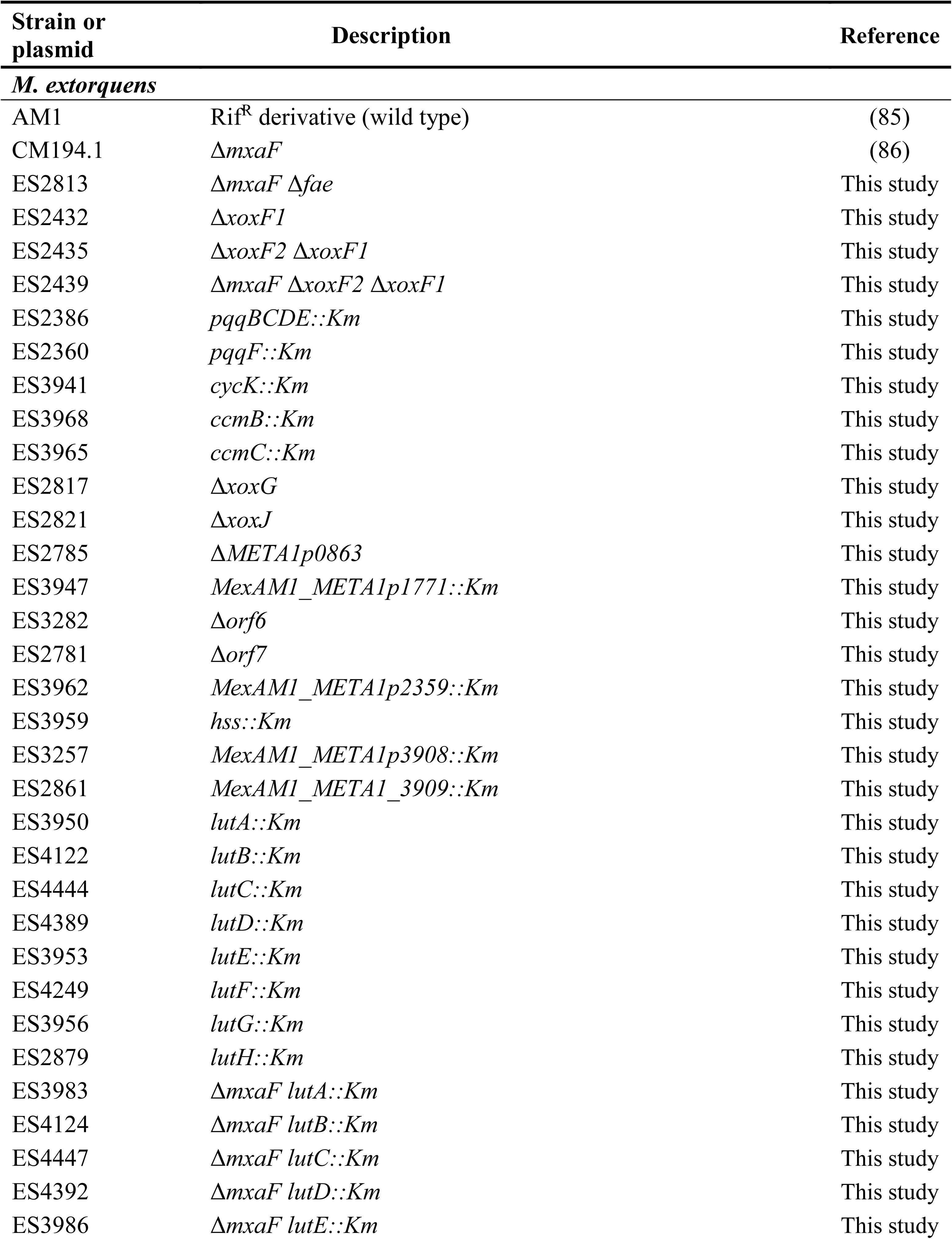
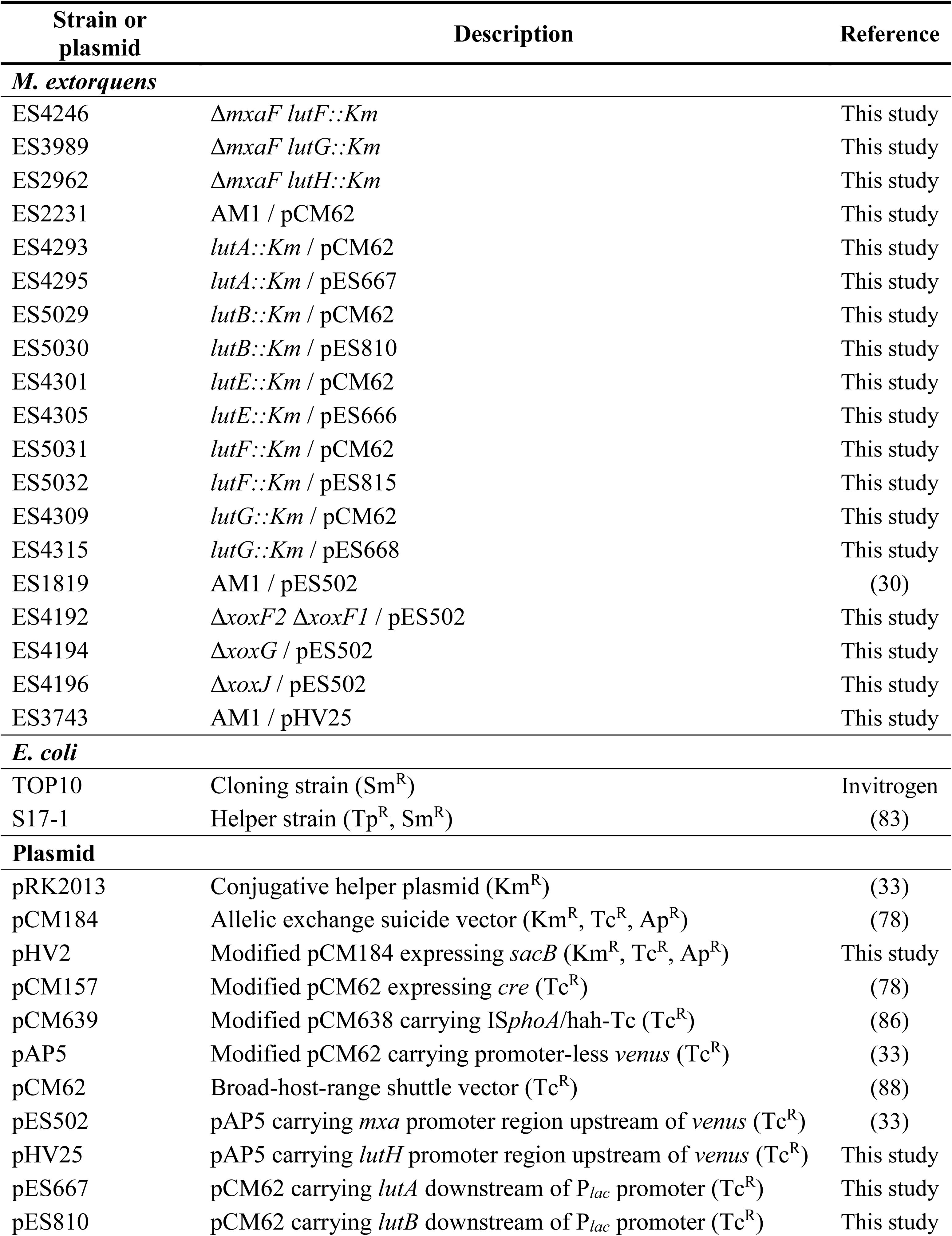
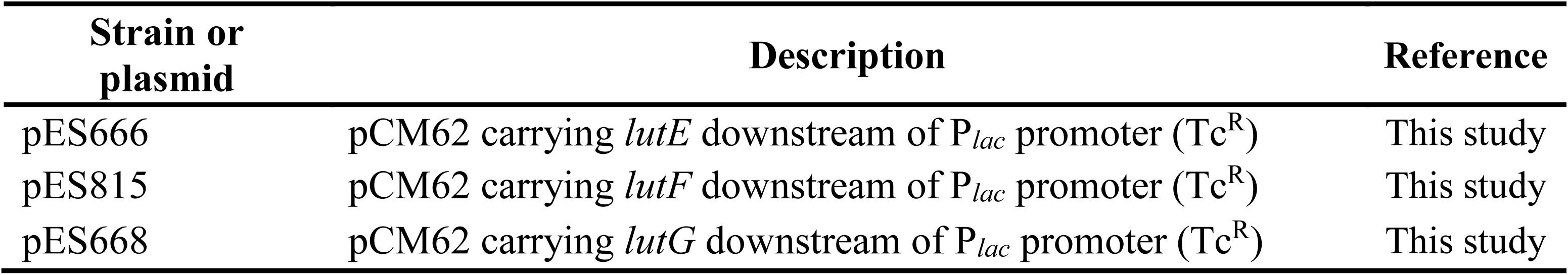
Strains and plasmids used in this study.

### Plasmid and strain construction

Primers used for plasmid and strain construction are listed in Table 6. The allelic exchange plasmid pHV2 was constructed by cloning the *sacB* gene from pCM433 (77) into the PscI site of pCM184 (78) in the same orientation as the Tc resistance gene. Insertion and orientation of *sacB* was confirmed by colony PCR. The *lutH* transcriptional reporter fusion was constructed by cloning the promoter region of *lutH* into the AclI and EcoRI sites upstream of a promoter-less *venus* gene in pAP5 (33). To create overexpression constructs for complementation studies, individual genes in the *lut* operon (*lutA*, *lutB*, *lutE*, *lutF*, and *lutG)* were cloned into the KpnI and SacI sites downstream of a P*_lac_* promoter in pCM62 (46). Diagnostic PCR was used to confirm successful integration of inserts. Plasmids were maintained in *E. coli* TOP10 (Invitrogen, Carlsbad, CA).

**Table 6.**
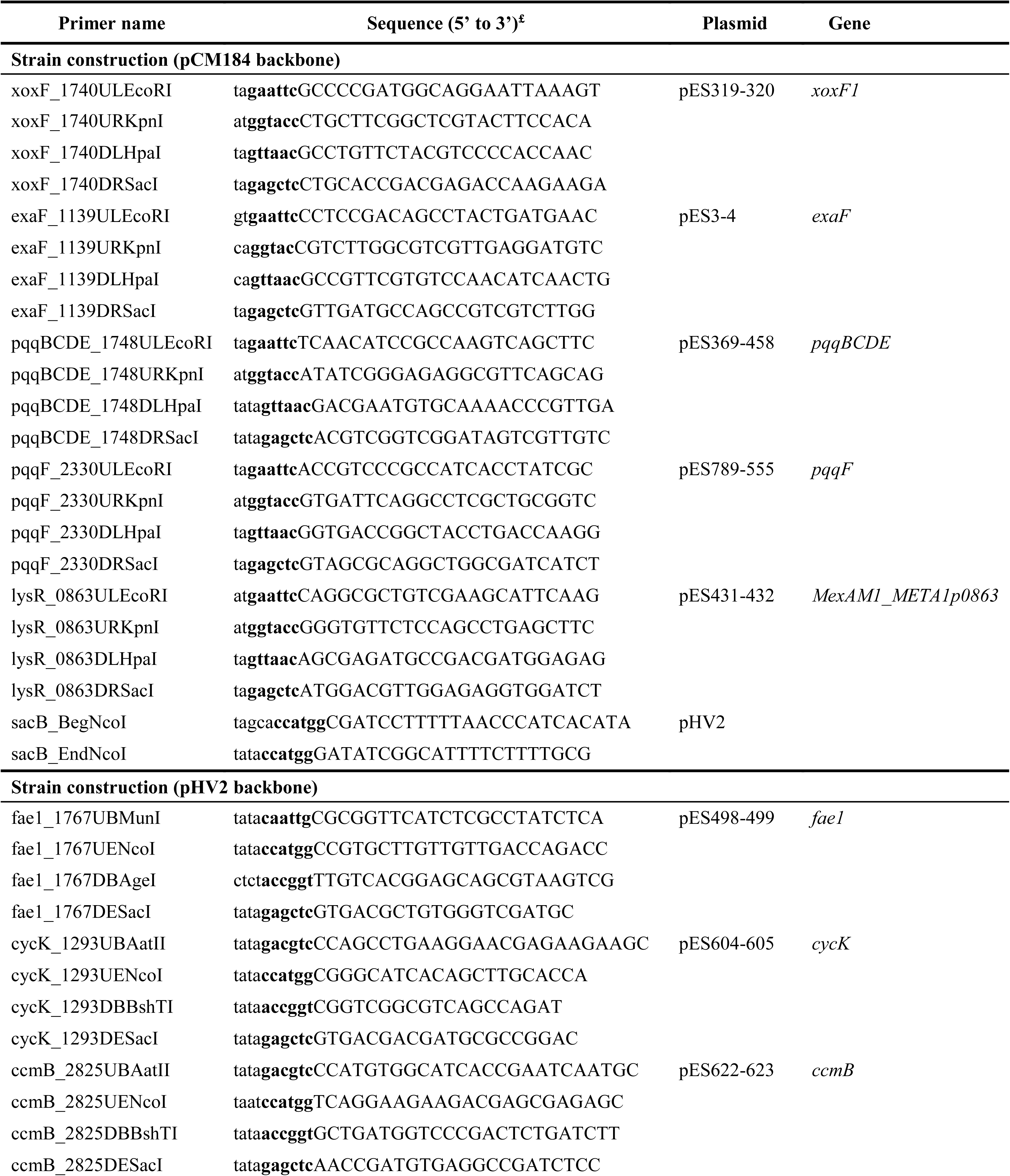
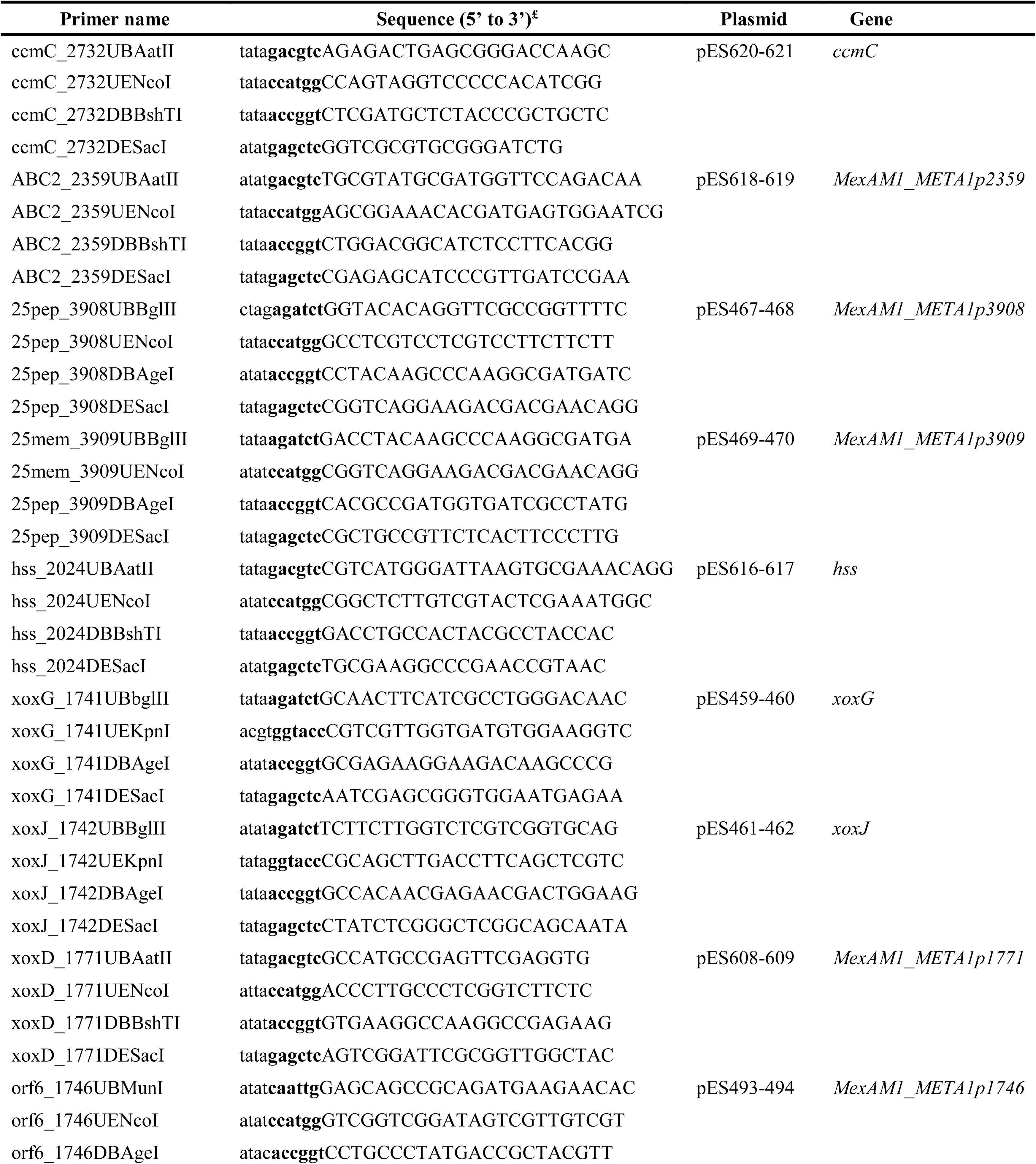
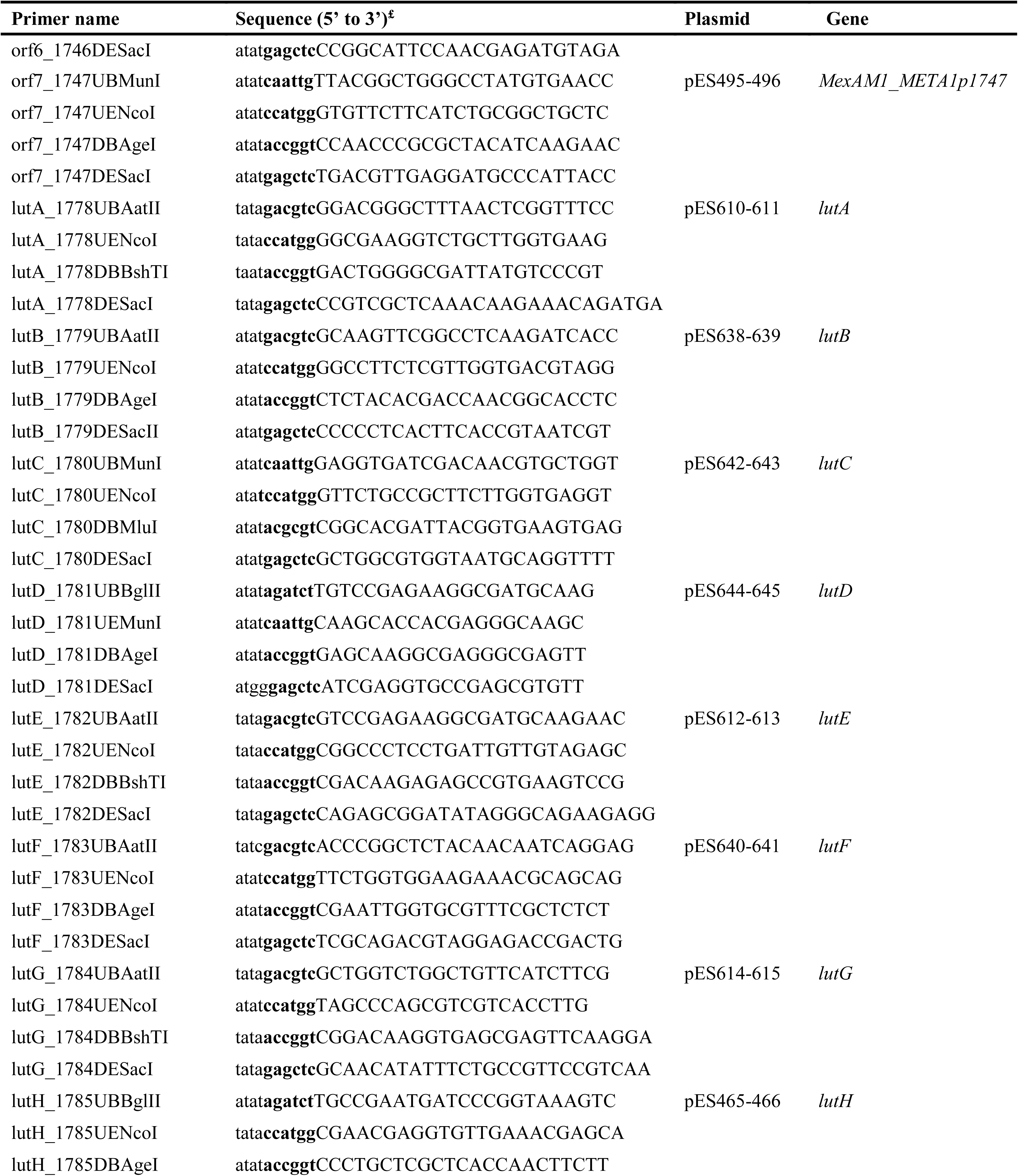
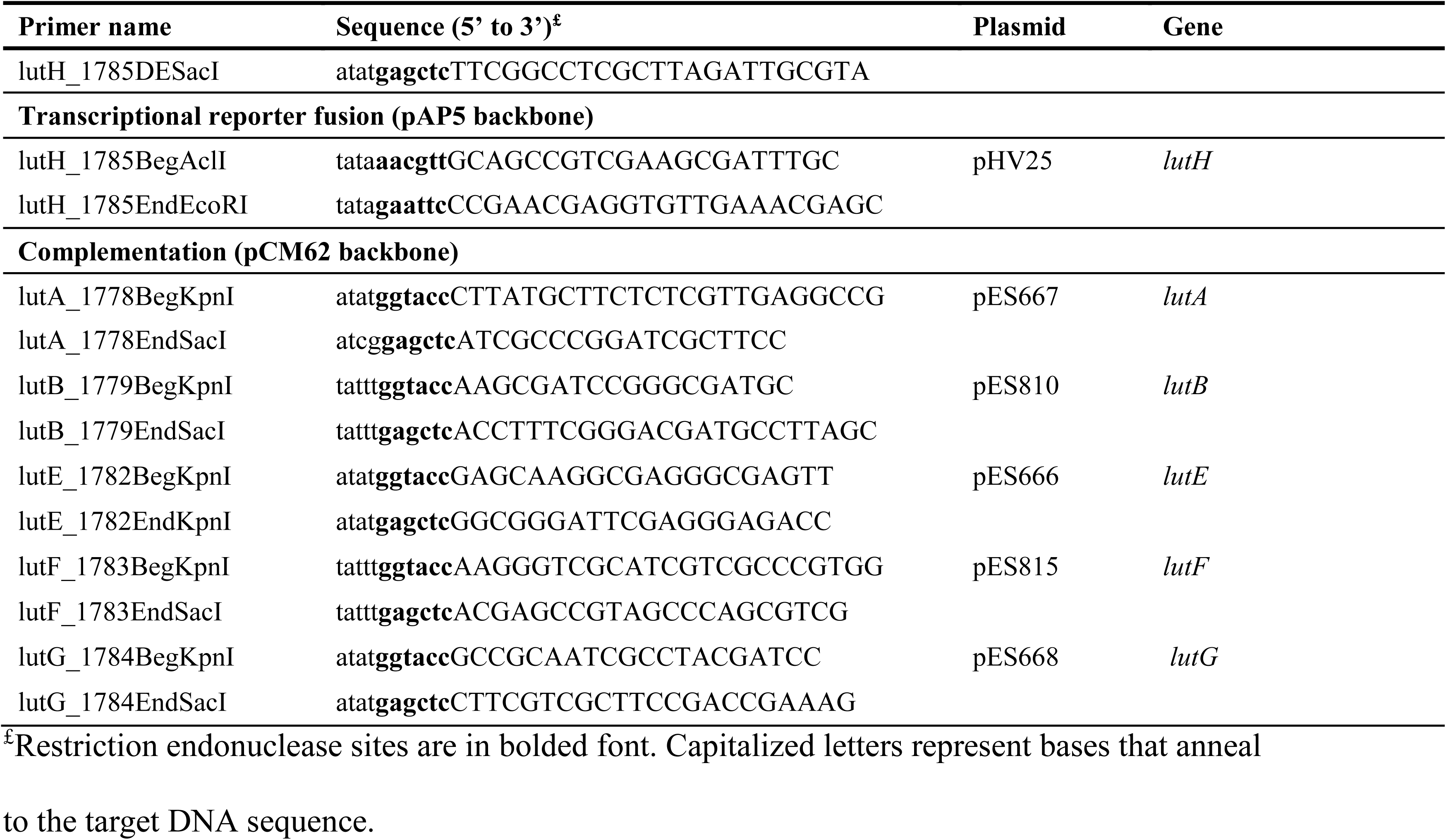
Primers used in this study.

Gene deletions were constructed using pCM184 or pHV2 as previously described (31) except 5% sucrose was added for counter selection against single crossovers (79) when using pHV2. Plasmids were conjugated into *M. extorquens* AM1 via biparental mating using *E. coli* S17-1 (81) or triparental mating using *E. coli* TOP10 (Invitrogen, Carlsbad, CA) and *E. coli* harboring the conjugative plasmid pRK2013 as described (31). When indicated, the Km resistance cassette was resolved using pCM157 to achieve marker-less deletions (80).

### Transposon mutagenesis

Suicide vector pCM639 carrying a mini transposon IS*phoA*/hah-Tc (82) was conjugated into the *mxaF fae* strain background via triparental mating as described (31, 83). Dilutions of the mating mixtures were plated onto MP succinate (15 mM) plus methanol (50 mM) La^3+^ medium containing 10 µg/mL Tc to select for successful integration of the mini transposon into the *M. extorquens* AM1 genome and 50 µg/mL Rif to counter select against *E. coli* strains bearing pCM639 or pRK2013. Plates were incubated for 5-7 days at 29°C. Transposon mutant colonies were streaked onto MP succinate methanol La^3+^ Tc medium for downstream studies.

### Location of transposon insertions

To identify the transposon insertion sites, genomic DNA was isolated using a Qiagen DNeasy UltraClean Microbial Kit (Qiagen, Germantown, MD). Degenerate nested PCR was performed as described (81, 82) with the following exceptions: PCR reactions contained 1 µM of each primer, 0.05 U/µL Dream Taq (Thermo Fisher Scientific, Waltham, MA), and 5% dimethyl sulfoxide. Modifications to the PCR amplification parameters included 2 minutes for the initial denaturation at 95°C, 6 cycles of annealing at 40°C followed by 25 cycles of annealing at 65°C for the first PCR reaction, and 30 cycles of annealing at 65°C for the second PCR reaction. PCR products were purified using a Qiagen QIAquick 96 PCR Purification Kit (Germantown, MD). Sequence analysis was performed using TransMapper, a Python-based program developed in-house to identify transposon insertion locations and map them to the *M. extorquens* AM1 genome for visualization using SnapGene Viewer (GSL Biotech LLC, Chicago, IL).

### Phenotypic analyses

Growth phenotypes were determined on solid or in liquid MP media using a minimum of three biological replicates. On solid media, colony size was scored after four days. Growth curve experiments were conducted at 29°C in an Excella E25 shaking incubator (New Brunswick Scientific, Edison, NJ) using a custom-built angled tube rack holder as previously described (26). Optical density (OD_600_) was measured at 600 nm using a Spectronic 20D spectrophotometer (Milton Roy Company, Warminster, PA). For strains with extended growth lags, suppression and acclimation was assessed. Strains from the growth curves were streaked onto methanol La^3+^ medium after they reached stationary phase. If the parent stock strain did not grow on methanol La^3+^ medium and the strain post-growth curve grew, acclimation versus suppression was tested. Strains were passaged from the methanol medium plate to a succinate medium plate. After colonies grew on succinate medium, they were streaked back onto methanol La^3+^ medium. If strains retained the ability to grow on methanol medium, it was concluded that growth was due to a suppressor mutation. If strains lost the ability to grow on methanol medium after succinate passage, it was concluded growth was due to acclimation and not a genetic change.

For Ln storage growth experiments, *mxaF* deletion strains were grown with 20 µM LaCl_3_ until six hours after transition into stationary phase. Cells were centrifuged and washed four times with MP medium lacking La^3+^ and a carbon source, and sub-cultured into fresh MP methanol medium without La^3+^. Six hours post stationary phase, the cultures were streaked onto MP methanol medium with and without La^3+^ to check for contamination. Cultures were again washed and sub-cultured as described above. This process was repeated for two rounds of growth without La^3+^.

### Transcriptional reporter fusion assays

*M. extorquens* AM1 strains carrying *mxa*, *xox1*, and *lutH* transcriptional reporter fusions (Table 5) which use *venus* (83) as a fluorescent reporter were grown in MP media supplemented with methanol only or methanol and succinate with and without La^3+^ as indicated in the text. Expression was measured as relative fluorescent units (RFU) using a SpectraMax M2 plate reader (Molecular Devices, Sunnyvale, CA) and normalized to OD_600_ as previously described (27).

### La^3+^ depletion during *M. extorquens* AM1 growth

Overnight cultures of wild type, *lutA*, *lutE*, *lutF*, and *lutH* mutant strains were inoculated 1:50 into 250 mL polycarbonate flasks (Corning Inc., Corning, NY) containing 75 mL of MP medium (76). Succinate (3.75 mM) and methanol (125 mM) were added as carbon sources with 2 μM LaCl_3_. Flasks were incubated at 28°C at 200 rpm in Innova 2300 shaking incubators (Eppendorf, Hauppauge, NY) for 44 h. To monitor La^3+^ depletion during *M. extorquens* AM1 cultivation, the Arsenazo III assay was used (84). 5 mL samples were collected at four different time points (OD_600_ of 0.04, 0.4, 0.7, and 1.5) and the concentration of La^3+^ remaining in the supernatant was calculated using the calibration curve prepared as previously described (84). A control of three uninoculated flasks containing MP medium with 2 μM LaCl_3_ were considered to determine La^3+^ adsorption by the flasks which was subtracted from the culture measurements. The initial concentration of La^3+^ in the media (before growth) was measured using the Arsenazo III assay in the same way as described above. Significant differences between depletion of La^3+^ by different strains were calculated using One-way ANOVA followed by a T-test.

### Cellular locations of Ln visualized using Transmission Electron Microscopy (TEM)

Sample preparation for TEM: wild type, *lutA*, *lutE*, *lutF*, and *lutH* mutant strains were grown in MP medium containing 125 mM methanol and 3.75 mM succinate as carbon sources with or without the addition of 20 μM LaCl_3_ until they reached an OD_600_ of ∼0.6. 3 mL of cells was harvested by centrifugation for 3 min at 1500 × *g* at room temperature and fixed for 30 min in 1 mL of 2.5% (v/v) glutaraldehyde (Electron Microscopy Sciences, Hatfield, PA) in 0.1 M cacodylate buffer (Electron Microscopy Sciences, Hatfield, PA). After fixation, cells were pelleted by centrifugation for 3 min at 1500 × *g* and washed with 1 mL of 0.1 M cacodylate buffer. Cell pellets were embedded in 2% (w/v) agarose and washed three times with 0.1 M cacodylate buffer. When indicated, pellets in agarose blocks were stained for 30 min in 1% osmium tetroxide in 0.1 M cacodylate buffer. Samples were washed three times with 0.1 M cacodylate buffer, dehydrated in acetone, and embedded in Spurr resin (Electron Microscopy Sciences, Hatfield, PA). Blocks were polymerized at 60°C for 48 h. 70 nm sections were obtained with a Power Tome XL ultramicrotome (RMC Boeckeler Instruments, Tucson AZ), deposited on 200 mesh carbon coated grids, and stained with 2% uranyl acetate (Electron Microscopy Sciences, Hatfield, PA). To assess the presence of La^3+^ by EDS, sections were left unstained. To image the distribution of cellular La^3+^, a TEM JOEL 1400 Flash (Japan Electron Optics Laboratory, Tokyo, Japan) was used. Detection of La^3+^ in the cells and high-resolution imaging were done with a JEOL 2200FS (Japan Electron Optics Laboratory, Tokyo, Japan) operated at 200kV. X-ray energy dispersive spectroscopy was performed using an Oxford Instruments INCA system (Abingdon, United Kingdom).

## ACKNOWLEDGMENTS

We would like to thank Dr. Lena Daumann and Dr. Nathan Good for critical review of this manuscript. We would like to thank San José State University General Microbiology students who isolated transposon mutants and purified genomic DNA for sequencing as part of a class research project. The mutant hunt in this study was inspired by Dr. Elizabeth Skovran’s undergraduate research mentor, Dr. Marc Rott at the University of Wisconsin-LaCrosse who used transposon mutagenesis in the teaching lab to identify genes involved in butanol oxidation in *Rhodobacter sphaeroides.* We would like to thank Timothy Andriese for assistance with sequencing of the transposon mutant DNA and all Skovran lab members for assistance with growth curves and transcriptional reporter fusion assays. TEM work was done at the Center for Advanced Microscopy, Michigan State University. We would like to thank Dr. Alicia Withrow for invaluable assistance with TEM experiments and Dr. Xudong Fan for his expertise using TEM-EDS. E.S., N.C.M-G., P.R-J., and H.N.V. directed experiments. E.S., N.C.M-G., P.R-J., and H.N.V. wrote the manuscript. P.R-J. conducted microscopy and Ln transport experiments. H.N.V, G.A.S, J.C., and E.C carried out transposon mutagenesis studies. H.N.V., G.A.S., R.C., J.C., C.R., E.M.A., E.C., N.F.L., and F.Y. constructed strains and conducted growth experiments. R.C., C.H., J.P.W., and G.A.S. conducted expression studies. R.T.N created TransMapper.

