## Supplemental Information for "Lanthanide transport, storage, and beyond: gene products and processes contributing to lanthanide and methanol metabolism in *Methylorubrum extorquens* AM1"

Supplemental Materials

**Fig S1. Complementation studies for the *lut* mutant strains.** Graphs depict representative data from biological triplicates grown in methanol media (125 mM) containing 2 µM $LaCl_3$ and 5 µg/mL Tc. Filled circles represent each mutant strain carrying the pCM62 empty vector. Empty circles represent each mutant strain with the respective gene cloned into pCM62.

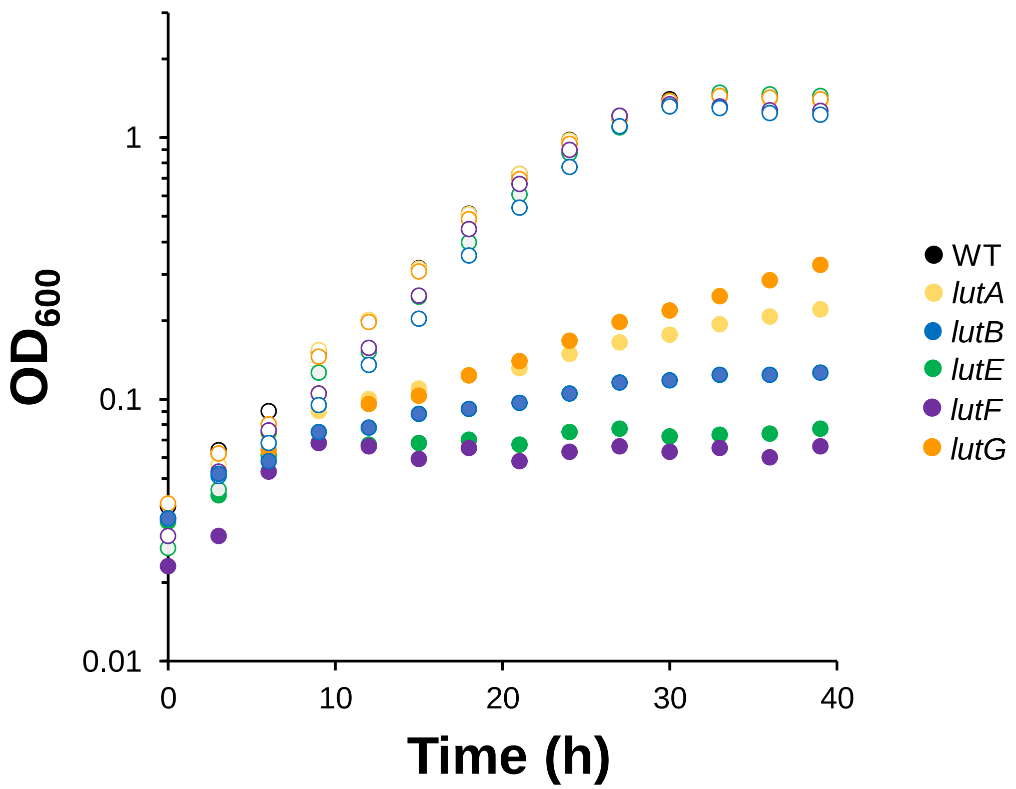

**Fig S2**. **Elemental analysis of cytoplasmic deposits.** Whole spectra of energy-dispersive X-ray analysis of the electron-dense deposits observed in the (A) cytoplasm of the wild-type (WT) strain grown with 20 µM $LaCl_3$ (B) cytoplasm of the wild-type strain grown without $LaCl_3$ and (C) periplasm of the *lutA* mutant strain grown with 20 µM $LaCl_3$. Microscopic images of these samples are depicted in Fig. 6 panel C and G and Fig. 4 panel F, respectively. Copper, carbon and silicon ion content is a result of these metals being present in the grids.

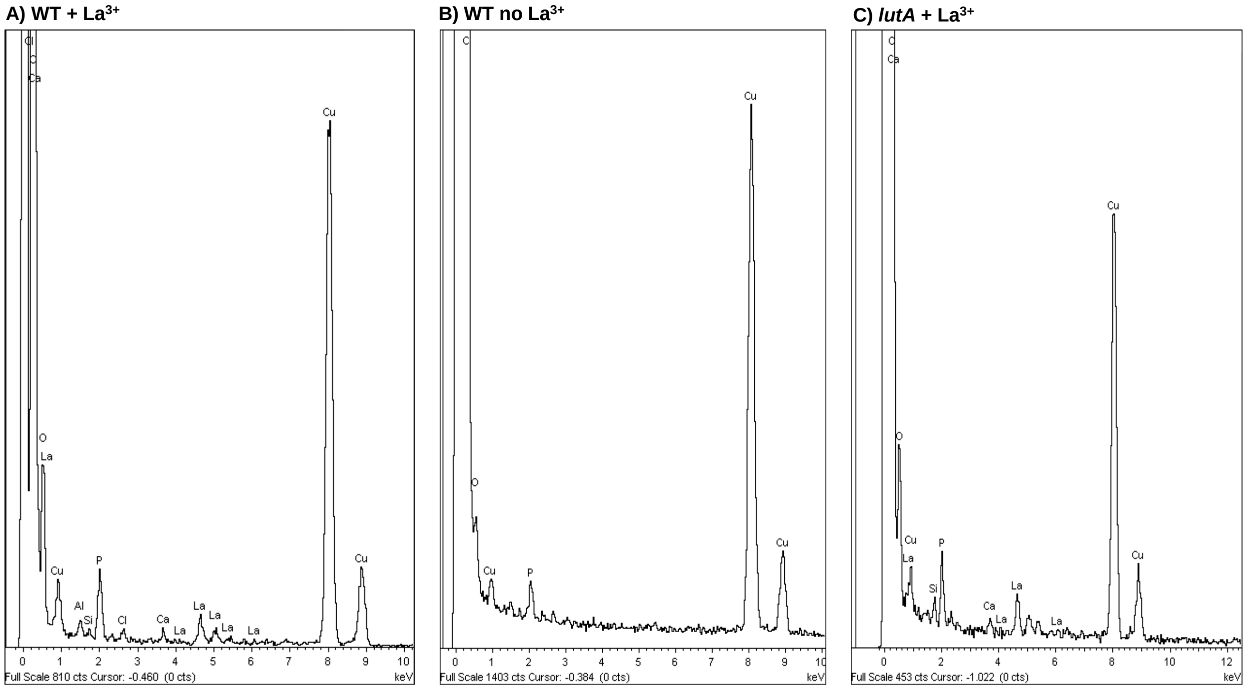
